# Engineering Plasmids with Synthetic Origins of Replication

**DOI:** 10.1101/2025.02.21.639468

**Authors:** Baiyang Liu, Xiao Peng, Matthew R. Bennett, Matthew R. Lakin, James Chappell

## Abstract

Plasmids remain by far the most common medium for delivering engineered DNA to microorganisms. However, the reliance on natural plasmid replication mechanisms limits their tunability, compatibility, and modularity. Here we refactor the natural pMB1 origin and create plasmids with customizable copy numbers by tuning refactored components. We then create compatible origins that use synthetic RNA regulators to implement independent copy control. We further demonstrate that the synthetic origin of replication (SynORI) can be engineered modularly to respond to various signals, allowing for multiplexed copy-based reporting of environmental signals. Lastly, a library of 6 orthogonal SynORI plasmids is created and co-maintained in E. coli for a week. This work establishes the feasibility of creating plasmids with SynORI that can serve as a new biotechnology for synthetic biology.

## Introduction

DNA plasmids play a vital role as mobile genetic elements in bacterial evolution, capable of mobilizing key functions such as antibiotic resistance^1–3^, resistance to heat^4^, and substrate metabolism^5–7^. Plasmids also remain the most common way to deliver recombinant DNA to a broad range of microbes^8–10^. However, most plasmids used today are limited to a few well-characterized types from the 1980s that have undergone minimal engineering, particularly in their origins of replication (ORI) which dictate plasmid compatibility and copy number^11,12^. As a result, researchers are largely restricted to a limited menu of plasmids that have fixed copy numbers and limited orthogonality, posing challenges for the advancement of plasmid-based applications such as implementing division of labor on plasmid level. In part, this lack of engineered plasmids is due to the non-modular genetic architectures of ORIs, which have evolved to incorporate features such as overlapping genes and operons. This lack of tunability, orthogonal variants, and modularity is in stark contrast to other genetic parts (e.g., promoters, RBS, terminators) that have been extensively engineered^13–15^.

To create synthetic plasmids beyond the bounds of evolution, we focused on creating a synthetic ORI (SynORI) that would address the current limitations of modularity, tunability, and orthogonality. To do this, we refactored and reengineered the pMB1 origin of replication (Fig. 1A). pMB1 belongs to the ColE1 family that initiates DNA replication through transcription and processing of an RNA primer (RNAII). DNA replication initiation is regulated by an antisense RNA (RNAI) that binds to the RNA primer co-transcriptionally to induce an alternative confirmation that terminates DNA replication^16^. As the concentration of antisense RNA is directly linked to plasmid copy number, this establishes negative feedback control on the replication of pMB1 plasmid.

**Fig. 1.**
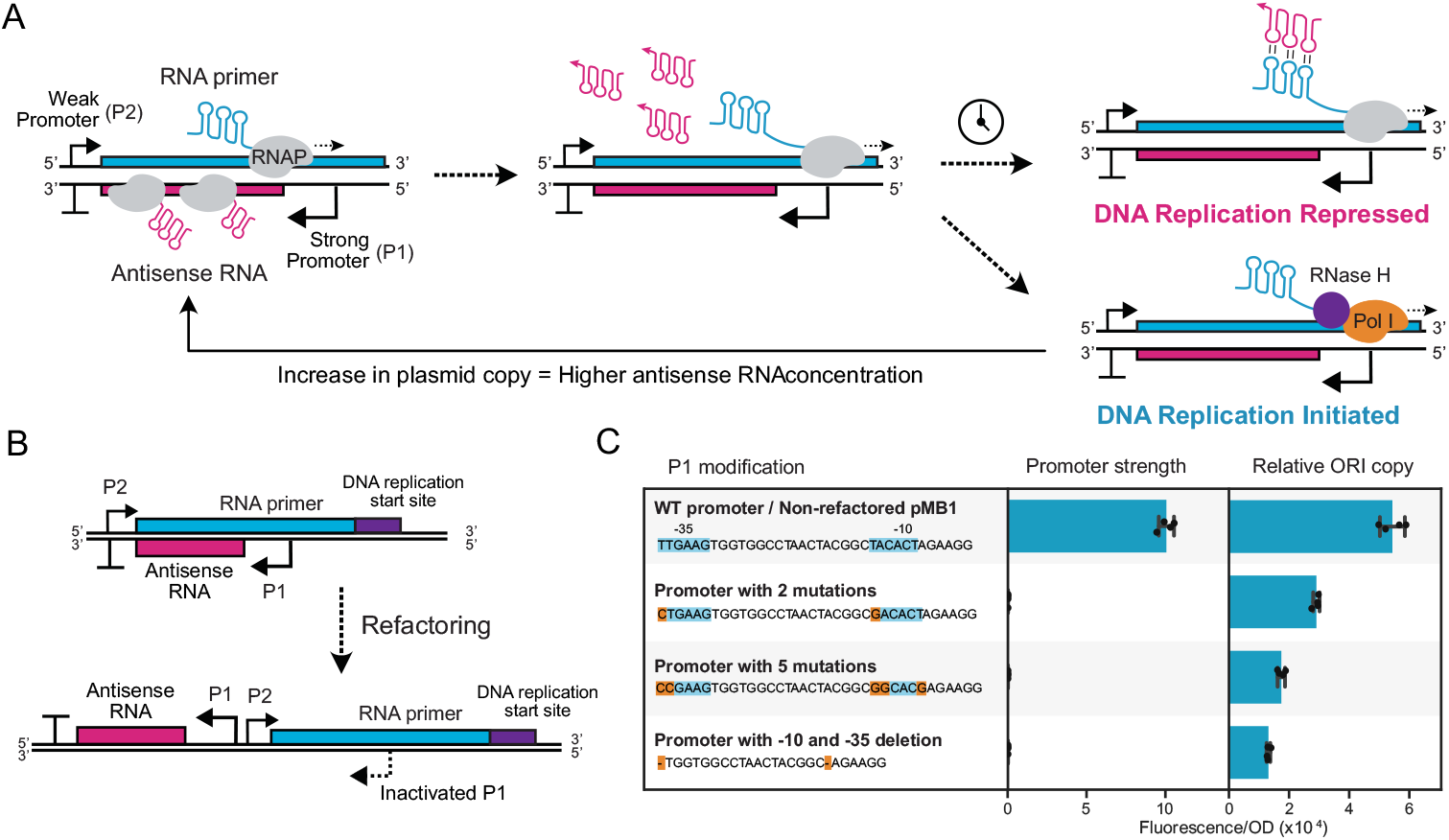
Refactoring the pMB1 origin of replication. (**A**) Schematic of pMB1 plasmid origin of replication (ORI) mechanism. The RNA primer is transcribed by RNA polymerase (RNAP) and by default folds into an RNA structure that is processed by RNase H to create a substrate for DNA polymerase I (Pol I) to initiate DNA replication. The antisense RNA is produced from an overlapping gene, which if allowed to interact with the RNA primer, induces an alternative fold that does not initiate DNA replication. As the concentration of antisense RNA is proportional to the plasmid copy, this serves as a copy-control negative feedback loop. (**B**) Schematic of a refactored pMB1 ORI that separates the RNA primer and antisense RNA gene and introduces inactivating mutations in the P1 promoter sequence. (**C**) Shows sequences of mutations to inactive the P1 promoter encoded inside the RNA primer. (Left graph) Whole-cell fluorescent characterization of the mutant promoters using a fluorescent reporter gene. (Right graph) Relative copy number of plasmids containing the original pMB1 ORI or refactored ORI with an RNA primer containing mutant P1 promoters. Copy number was characterized by encoding a constitutive RFP expression cassette onto the plasmid. Fluorescence characterization was performed (measured in units of fluorescence/optical density (OD) at 600 nm) in *E. coli* cells. Data show mean and individual values of n = 4 biological replicates.

### The pMB1 ORI can be refactored

To refactor the overlapping components of pMB1 ORI, we evaluated whether the antisense RNA promoter inside the RNA primer, called P1, could be removed without disrupting the replication capability of the ORI (Fig. 1B). A collection of P1 promoters mutated in their -10 and -35 regions were designed and their relative strength measured in *E. coli* cells using RFP expression (Fig. 1C and fig. S1). No promoter activity was observed from any of the modified promoters. We then built a refactored pMB1 ORI by separating the antisense RNA and RNA primer into separate transcription cassettes and using an RNA primer that contained mutations in the P1 promoter. To measure the relative copy number, a constitutively expressed RFP was inserted into each refactored plasmid, and whole-cell fluorescence measured. A reduction in the relative copy number was seen with greater modification in the RNA primer (Fig. 1C), with the 2-mutation promoter having the closest copy number to the naïve pMB1 ORI, which was used for subsequent iterations. Taken together, these results show we can effectively refactor the pMB1 ORI by separating transcription cassettes and engineering the RNA primer.

### Synthetic ORI (SynORI) can be created using synthetic transcriptional repressors

We next sought to create a synthetic origin of replication (SynORI) in which the natural pMB1 ORI copy control mechanism was replaced with synthetic transcriptional repressors. Specifically, we focused on the pT181 transcriptional attenuator, a transcriptional regulator discovered in a naturally occurring plasmid from *Staphylococcus aureus*^17^, which has been extensively engineered to yield orthogonal libraries^18,19^. This system is composed of a target RNA that harbors a latent transcriptional terminator that adopts a conformation that prevents downstream transcription only in the presence of its corresponding repressor small RNA (sRNA) (fig. S2). We reasoned that, by inserting the RNA primer downstream of the target and replacing the antisense RNA with the repressor sRNA, we could effectively implement negative feedback to control the plasmid copy number in SynORI (Fig. 2A). By separating copy-control and replication initiation into distinct, interchangeable genetic modules, this method offers enhanced design flexibility. To test this hypothesis, a set of SynORIs using a pair of orthogonal pT181 attenuators that contained all four combinations of target and repressor sRNAs was created and the relative copy number characterized. Designs with cognate pairs of target and repressor that can implement negative feedback showed a dramatic decrease in relative copy number in *E. coli* cells compared to designs using non-cognate pairs (Fig. 2B, fig. S3A). Interestingly, we saw the same trend in *Shewanella oneidensis*, suggesting that SynORI functions in multiple cell types (fig. S4). These results suggest that SynORIs can be created that implement plasmid copy control using synthetic transcriptional regulators.

**Fig. 2.**
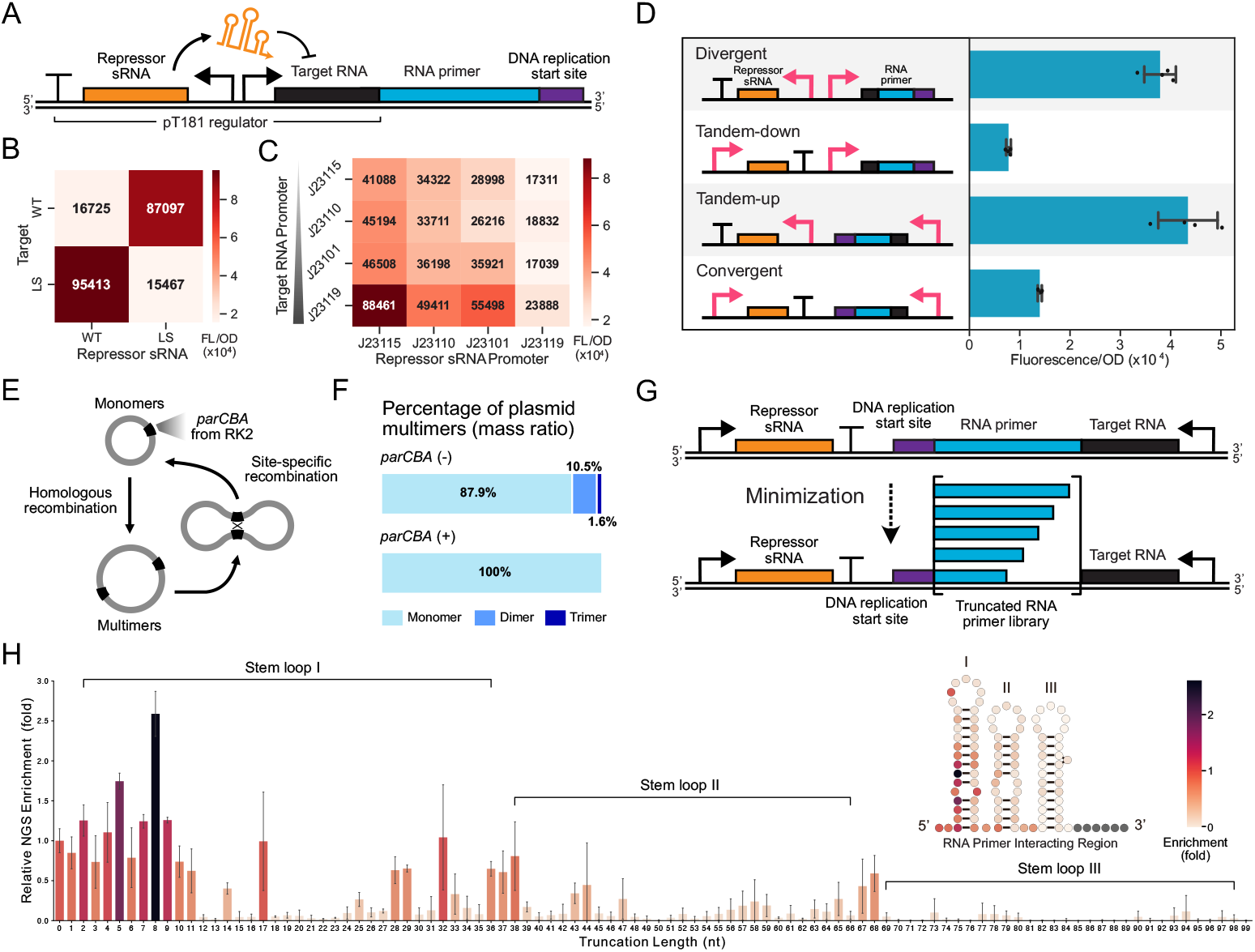
Creating a synthetic origin of replication (SynORI) through reengineering the pMB1 ORI. (**A**) Schematic of synthetic ORI (SynORI) in which the copy-control mechanism is replaced with the synthetic pT181 transcriptional attenuator. In this mechanism, the repressor small RNA (sRNA) inhibits transcription of a target RNA that is placed upstream of the RNA primer to control plasmid copy number. Matrix showing the relative copy number of plasmids containing (**B**) different combinations of target and repressor sRNA from two orthogonal pT181 variants (WT and LS) and (**C**) different strength promoters. (**D**) Relative copy number of plasmids with different SynORI genetic architectures. Relative copy number was characterized by encoding a constitutive RFP expression cassette onto the plasmid in *E. coli* cells (measured in units of Fluorescence/Optical density [OD] at 600 nm). (**E**) Schematic of multimer resolving by *parCBA*. (**F**) Percentage of plasmid multimers in *E. coli* cells with and without *parCBA* present. Data shows the mass ratio (percentage of total nucleotides in each multimer species) measured by nanopore whole-plasmid sequencing. (**G**) Schematic of the RNA primer truncation library. (**H**) Relative enrichment of each truncation (5’ to 3’) using a growth-based selection. Relative enrichment was determined by comparing the abundance of each design in a naïve library before the Golden Gate assembly to the abundance of each design after transformation, selection, and plasmid isolation. The enrichment of the plasmid variant without truncation (0 nt) was normalized to 1. Only truncations that preserve plasmid replication should be enriched following the selection. Insert shows the predicted structure of the RNA primer with each nucleotide’s color corresponding to the relative enrichment at that truncation site. In B, C, F data shows mean of n = 3 biological replicates and in D, H data shows values of n = 4 biological replicates.

To determine whether SynORI offers tunable copy control, we replaced the promoters driving both target and repressor sRNA with four varying strength promoters. From this, we saw the copy-number varied over 5-fold, with the highest copy number seen when target RNA transcription was greater than the repressor sRNA, and the lowest copy number when transcription rates were inverted (Fig. 2C, fig. S3B). Next, we investigated how the orientation of transcriptional cassettes affected the copy number (Fig. 2D). Tandem-down and convergent orientations showed the lowest plasmid copy, likely due to the repression of RNA primer transcription from DNA supercoiling created by the transcription of the upstream repressor sRNA^20,21^. Taken together, this shows SynORI copy number can be tuned using promoter libraries and distinct genetic architectures, which are unfeasible with the naïve pMB1 ORI.

Plasmid multimerization is a persistent problem seen in ColE1 plasmid family^22–24^, which in part stems from the deletion of the recombination site, *cer*, from the laboratory derivatives that resolve multimers^25^. To investigate multimerization in our SynORI, we sequenced DNA plasmids from *E. coli* cells using whole-plasmid sequencing. We observed ∼10.5% and ∼1.5% of plasmids formed dimers and trimers, respectively. Surprisingly, reintroducing *cer* into SynORI did not reduce multimer formation (fig. S5). However, incorporating a parCBA system, adapted from plasmid RK2^26^ (Fig. 2E), effectively eliminated multimers (Fig. 2F). The addition of a toxin-antitoxin system, parDE, further enhanced plasmid stability^4^. These findings demonstrate that natural plasmid maintenance mechanisms can effectively mitigate SynORI multimerization.

As SynORI is independent of the naïve RNA primer’s copy control mechanism, we reasoned that the RNA primer could be minimized. Deletion of the entire region responsible for interacting with the antisense RNA (termed stem loop I, II, and III) proved non-functional in *E. coli*. Therefore, we conducted incremental truncations from the 5’ end of the RNA primer using a pooled library approach (Fig. 2G). By sequencing and analyzing the enrichment of various truncations after transformation into *E. coli*, we found that approximately two-thirds of the interacting sequence (stem loop I and II) could be removed without compromising functionality (Fig. 2H, fig. S6). However, truncations inside stem loop III led to non-functional SynORIs, which we verified by individual plasmid characterization (fig. S7). Interestingly, truncations within stem loop structures often disrupted RNA primer function, suggesting that an incomplete stem loop disrupted RNA primer function. Taken together, these results show that a functional and minimal SynORI design can be achieved using an RNA primer truncated to stem loop III, in combination with a tunable synthetic promoter, a convergent genetic architecture, and a parCBA resolvase system.

### Benchmarking the performance of pSynORI

To evaluate the overall performance of the finalized SynORI plasmids (pSynORI) we compared it with pMB1 in *E. coli*, with regard to several important performance criteria. Cells containing pSynORI had generally comparable growth curves to cells containing pMB1, although maximum growth was slightly reduced (Fig. 3A). The plasmid copy distribution, measured using single-cell fluorescence, showed a slightly wider copy distribution for pSynORI (Fig. 3B), likely due to the intrinsic difference in the pMB1 and pSynORI copy control mechanisms. Next, we evaluated the rate of multimer formation. As expected, multimers were not observed for pSynORI; however, significant levels of multimers (dimers, timers, and tetramers) were seen for pMB1. Plasmid copy number stability was next considered by measuring whole-cell fluorescence over a 7-day continuous culture with antibiotic selection. Both pMB1 and pSynORI exhibited relatively stable whole-cell fluorescence levels over 7 days (Fig. 3D). Finally, we quantified the rate of spontaneous plasmids loss by comparing colony forming units (CFU) with and without antibiotic selection of plasmids after overnight growth (Fig. 3E). No plasmid loss was observed in cells with pMB1 and pSynORI in both culture conditions. Taken together, these results indicate that pSynORI performs comparably to the original pMB1 plasmid.

**Fig. 3.**
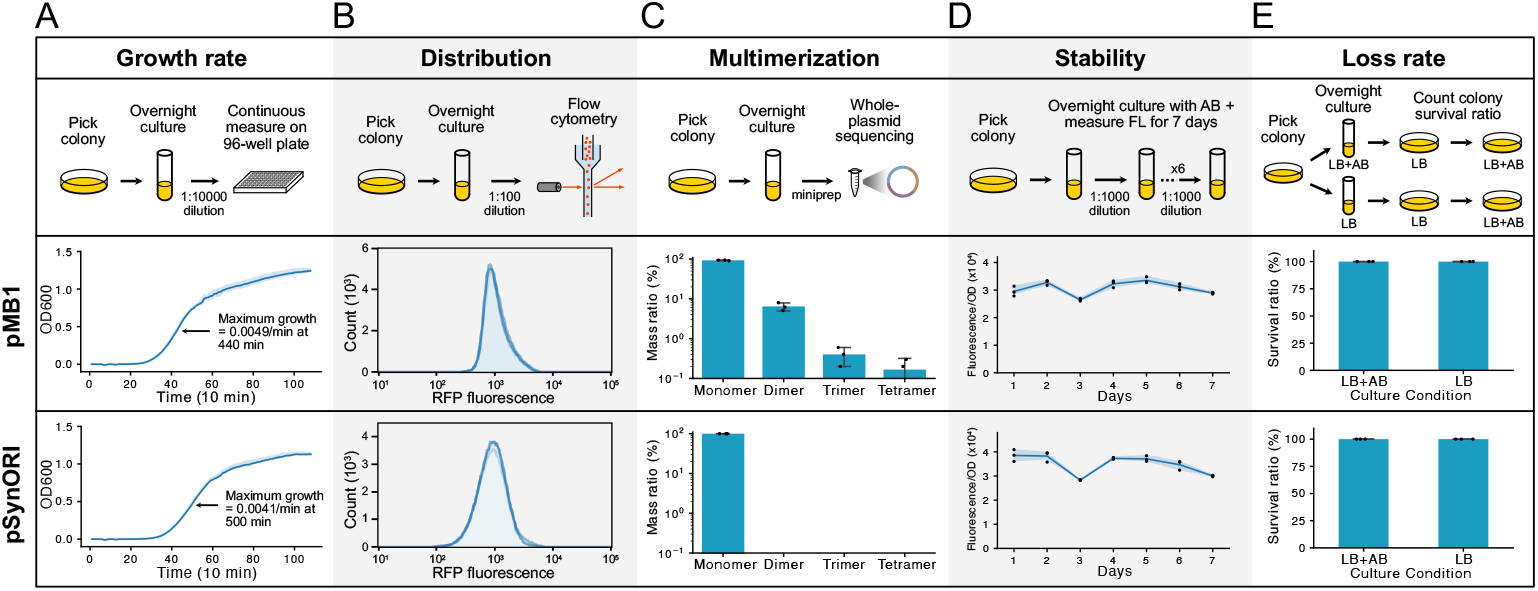
Benchmarking the performance of pMB1 and pSynORI. (**A**) Characterization of growth rate in *E. coli* cells transformed with pMB1 and pSynORI (measured in units of optical density [OD] at 600 nm). Maximum growth rate is indicated. (**B**) Single-cell fluorescence of *E. coli* cells transformed with pMB1 and pSynORI containing a constitutive RFP expression cassette. The coefficients of variation (CV) of 3 biological replicates of pMB1 are 56.9, 57.8, 57.9. The CVs of pSynORI are 60.1, 68.2, 60.0. (**C**) Percentage of plasmid multimers in *E. coli* cells with and without *parCBA* present. Data shows the mass ratio (percentage of total nucleotides in each multimer species) measured by nanopore whole-plasmid sequencing. (**D**) Relative copy number of plasmids over time. Relative copy number was characterized by encoding a constitutive RFP expression cassette onto the plasmid in *E. coli* cells (measured in units of Fluorescence/Optical density [OD]). (**E**) The percentage of cells retaining plasmid after culturing is measured by comparing the percentage of cells surviving with and without antibiotic (+AB) selection. Cells retaining plasmid are resistant to AB and should grow in both conditions. Data A-D shows n = 3 biological replicates. Data in E shows data from 20 colonies collected from n = 3 biological replicates.

### Converting chemical signals into plasmid copy

An advantage of the pSynORI system is its flexibility in adapting different regulatory systems for plasmid copy control. To demonstrate this, we explored ligand-inducible copy control using inducible promoters and transcriptional riboswitches (Fig. 4A). By using the IPTG- and cumate-inducible promoters to transcribe the pT181-regulated RNA primer, we achieved up to 13-fold control of copy number (Fig. 4, B and C). Additionally, we replaced the pT181 target RNA with the ZTP- and 2AP-riboswitches^27–29^, enabling both low-to-high and high-to-low plasmid copy flipping in response to specific chemical signals (Fig. 4, D and E). Furthermore, we demonstrated the integration of multiple signals for plasmid copy regulation by using the cumate- and IPTG-inducible promoters to control RNA primer and repressor sRNA transcription independently. As expected, plasmid copy was upregulated by cumate and downregulated by IPTG (Fig. 4F, fig. S8). These results highlight pSynORI’s potential as a flexible platform for converting chemical information into plasmid copy number, providing a novel reporting modality that can be easily analyzed through sequencing.

**Fig. 4.**
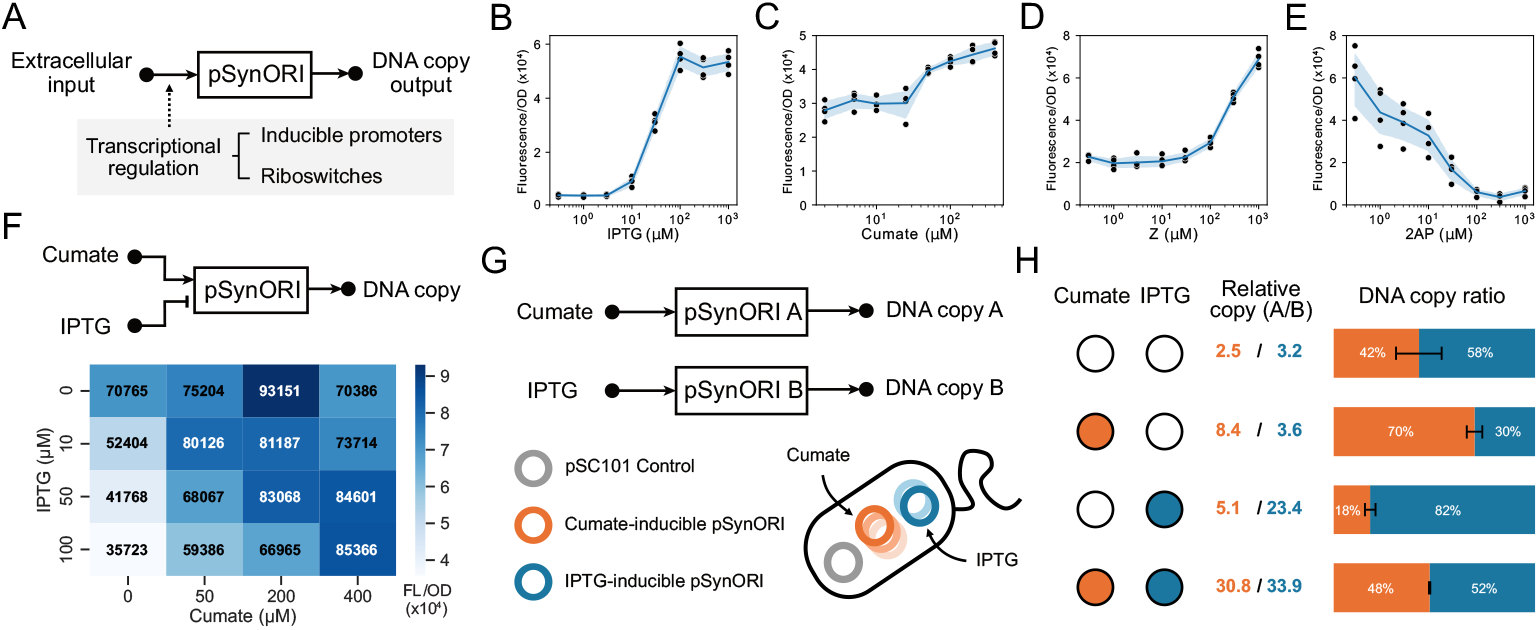
pSynORI provides a modular platform to transform chemical signals into DNA copy output. (**A**) SynORI plasmids (pSynORI) can be programmed to convert distinct chemical signals into DNA copy output using transcriptional regulators to control RNA primer transcription. Relationship between relative copy number and chemical input for pSynORI built using (**B**) Plac (activated by IPTG), (**C**) PcymR (activated by cumate), (**D**) the ZTP riboswitch (activated by Z), (**E**) and the yxjA riboswitch (repressed by 2AP). (**F**) Matrix showing the relative copy of a pSynORI that is activated by cumate and repressed by IPTG with inducer titrations. Relative copy number was characterized by encoding a constitutive RFP expression cassette onto the plasmid in *E. coli* cells (measured in units of Fluorescence/Optical density [OD] at 600 nm). (**G**) Schematic of orthogonal and inducible pSynORI that convert chemical input signals into DNA copy, which can be read using sequencing. A pSC101 plasmid with a non-inducible copy number is included as a reference. (**H**) The relative copy of pSynORI A and B was calculated using the reference plasmid. Bar plots show the DNA copy ratio of pSynORI A and B. Data in B-E and F shows n = 4 biological replicates and Data in H shows n = 3 biological replicates.

Next, we investigated the potential of pSynORI for multiplexable reporting of chemical signals through copy control. To test this concept, we constructed two orthogonal pSynORI (A and B) responding to cumate and IPTG, respectively (Fig. 4G). These pSynORI were co-transformed into *E. coli* cells alongside a reference plasmid, pSC101, that should maintain a steady copy number. Cells were exposed to different chemical combinations, and plasmid copy numbers were determined through whole plasmid sequencing (Fig. 4H). Relative to the reference plasmid, pSynORI A and B showed increases in copy number upon addition of their corresponding chemical signal, although the exact relative copy level achieved varied between conditions. The relative ratio of these plasmids also adjusted as expected, which was further validated using fluorescent reporters (fig. S9). Additionally, the dynamic responses of single-plasmid and double-plasmid inducible SynORI systems were characterized using a microfluidic device (fig. S10, Movie S1 and S2). These results show pSynORI can be used for multiplexing the reporting of chemical signals using easy-to-read plasmid copy numbers.

### Six orthogonal pSynORI co-exist in the same cell

An advantage of pSynORI is the ability to leverage orthogonal regulatory systems for plasmid copy control, enabling the creation of compatible plasmids that coexist within the same cell. To demonstrate this, we constructed seven pSynORI plasmids using previously described orthogonal pT181 systems (YS, F6, F4, F4m1, F3m1, F15m5, and LS)^19^. We initially validated the ability of different target and repressor sRNA pairs to regulate copy control in pSynORI. For each pT181 variant, a pSynORI with a matching target and repressor sRNA was compared with designs containing non-matching target or repressor sRNA from another variant, LS. All tested pT181 pairs showed clear repression of pSynORI copy only when the target and repressor sRNA were matched (Fig. 5A, fig. S11A). Interestingly, the final copy number of each pSynORI varied (3.4-fold), which we reasoned was due to the difference in repression efficiency of these pT181 variants^19^ or potential sequence-specific interactions with the downstream RNA primer. These results demonstrate a library of pSynORI plasmids can be created using orthogonal transcriptional regulator libraries.

**Fig. 5.**
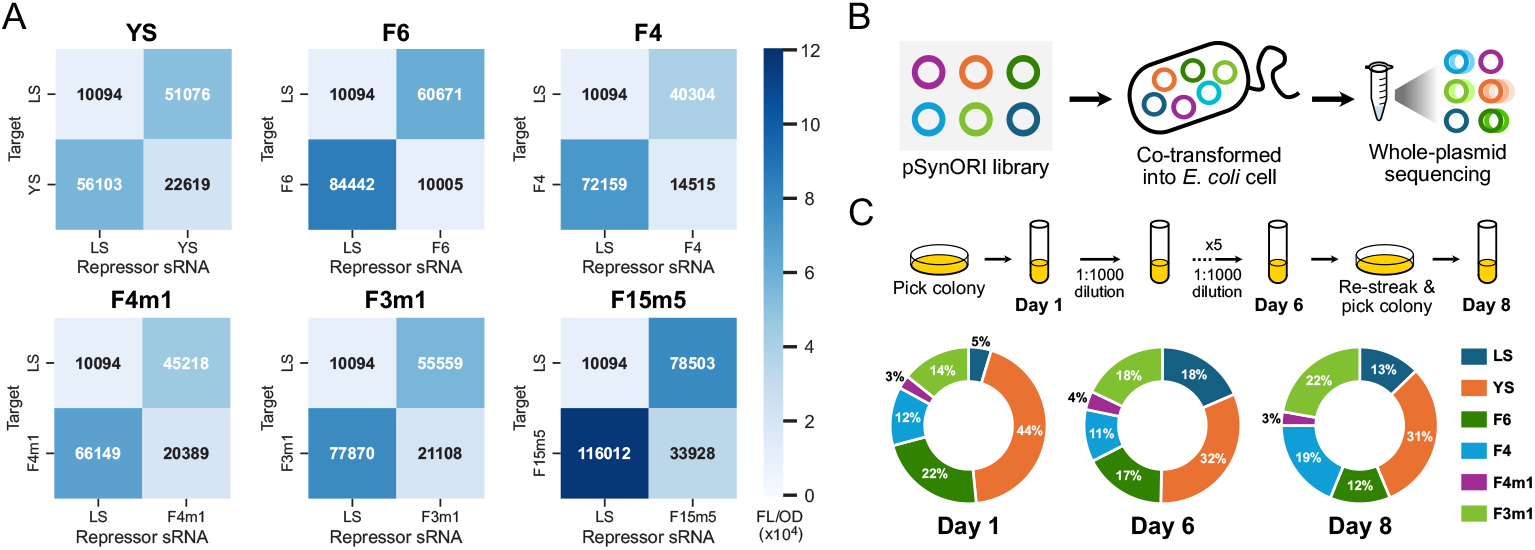
A library of orthogonal and compatible pSynORI. (**A**) Matrices showing the relative copy number of pSynORI built with 7 orthogonal pT181 variants. For each variant, combinations with a non-matching target and repressor sRNA (LS) are also included as a control. (**B**) A library of 6 pSynORI with different pT181 regulation systems (LS, YS, F6, F4, F4m1, F3m1) and antibiotic resistances (CmR, SpcR, KanR, AmpR, TetR, AprR) was created and co-transformed into *E. coli* cells. The proportions of plasmids in a cell population are measured using nanopore whole-plasmid sequencing. (**C**) Plasmid proportion during 8-day continuous culture with antibiotic selection. Cells are re-streaked on day 6, and individual colonies grown from a single cell were picked for day 8 cultures. The samples from day 1, 2, 4, 6, and 8 were processed for sequencing to confirm the presence of plasmids. The ring plot shows the ratio of plasmids on day 1, 6, and 8, with other days shown in figure S11. Data in A shows n = 4 biological replicates and data in C shows n = 3 biological replicates.

We then picked six SynORI variants that exhibit strong plasmid copy repression to create a library of compatible pSynORI. Each pSynORI was cloned to contain a distinct antibiotic resistance gene. Surprisingly, colonies were obtained from a single transformation of all six pSynORI into *E. coli* cells. To verify the presence of all six plasmids, we conducted a 6-day continuous culture with antibiotic selection. After 1, 2, 4, and 6 days, plasmids were extracted, and sequenced, and the presence of the six plasmids confirmed (Fig. 5C, fig. S11B). Although plasmid distribution was uneven, likely due to varying pT181 repression efficiencies and potential crosstalk, we observed no plasmid loss throughout the culture. Interestingly, a mutation in the LS-SynORI plasmid on day 4 in one biological repeat led to a change in plasmid copy from low to high (fig. S11B), suggesting that the antibiotic concentrations might require optimization based on plasmid composition and that maintaining six plasmids simultaneously can be challenging for the cell. As a control, we also constructed six pMB1 plasmids with distinct antibiotic-resistance genes. Attempts to transform the six pMB1 were unsuccessful (fig. S12A), although colonies were obtained by transforming four pMB1 (fig. S12B). A mixture of small and large colonies was observed and attempts to grow cultures resulted in no growth for some colonies. For cultures that grew, analysis of isolated DNA plasmids showed a high frequency of dimer formation (fig. S12C). Taken together, these results suggest our modifications to pSynORI are critical for achieving compatibility.

We next aimed to provide evidence that cultures were composed of cells containing all six plasmids, rather than a consortium of cells that had escaped individual plasmids. To test this, an aliquot of cells from the day 6 culture was plated and grown overnight. Single colonies grown from individual cells were then used to inoculate a culture grown until day 8 and the presence of all six plasmids was confirmed. In conclusion, we have successfully demonstrated the creation of a compatible library of six pSynORI that can co-exist in the same cell, which extends the current state of the art in plasmid design.

## Discussion

Our pSynORI system demonstrates that plasmids can be extensively refactored and modified to achieve desired plasmid features of tunability, orthogonality, and modularity in replication control. In contrast to previous works on engineering plasmid origins of replication^30,31^, pSynORI features refactored regulatory elements that effectively modularize plasmid copy regulation and replication initiation. This modular design of pSynORI creates flexibility by making it easier to add desired regulatory elements without unintentionally impairing replication. As such we anticipate pSynORI will open applications such as an ability to integrate diverse input signals into copy control for biosensing applications, synthetically coupling plasmid populations for dynamic adaption^32^ and signal transduction, dynamically controlling plasmid copy to decouple growth and production phases^33^, introducing a biocontainment measure to restrict the spread of plasmids^34^, serving as a platform for the division of labor at the plasmid level^35^, and enabling the development of modular and self-regulating metabolic pathways^36^.

While we focused here on refactoring and reengineering the pMB1 plasmid, we expect the approach to be broadly applicable to other plasmids^37^. This would be useful to apply to plasmids suitable for non-model microbes, which often lack plasmids that are functional, compatible, or cover diverse copy ranges. Applying this approach to extremely broad host range plasmids discovered within plasmidomes^38^, in combination with broad-host or conjugative genetic parts^8,39–,42^, could allow the creation of pSynORI adapted for programming diverse microbes or microbiomes. In summary, the development of pSynORI could play a vital role in establishing tunable, orthogonal, and modular plasmid-based synthetic biological systems.

## Methods

### Plasmid assembly strategies

All plasmids used in this study are listed in Table S1. The key plasmids are visualized in Fig. S13. The sequence of pSynORI plasmid characterized in Fig. 3 is provided in Table S2, and the sequences of promoters, riboswitches, and pT181 transcriptional repressors are provided in Table S3-5. The examples of RNA primer truncations used in Fig. 2G and 2H are provided in Table S6. Plasmids were constructed with a combination of inverse PCR, Gibson assembly, or Golden Gate assembly. The promoters and pT181 were replaced using a flexible design of Golden Gate assembly, which is explained in Table S7 with key primer sequences. All assembled plasmids were verified using Sanger DNA sequencing (Genewiz) or Nanopore sequencing (Plasmidsaurus).

### Plasmid transformation and culture conditions

For experiments using *E. coli*, plasmids were transformed into chemically competent *E. coli* (NEB Turbo strain). *E. coli* cells were recovered at 37 °C for 1 hour, plated on LB-agar plates containing combinations of 34 μg/mL chloramphenicol (Sigma-Aldrich), 50 μg/mL spectinomycin (Sigma-Aldrich), and 100 μg/mL kanamycin (Sigma-Aldrich) depending on the plasmids used, and incubated overnight at 37 °C. For the transformation of six plasmids, the LB-agar plates contained 34 μg/mL chloramphenicol, 50 μg/mL spectinomycin, 50 μg/mL kanamycin, 100 μg/mL carbenicillin (Sigma-Aldrich), 50 μg/mL apramycin (Sigma-Aldrich), and 5 μg/mL tetracycline (Sigma-Aldrich). 1.25 μg/mL tetracycline was used for liquid LB culture as tetracycline showed higher selection efficiency in liquid LB than on LB-agar plates.

For induction experiments using *E. coli*, colonies were picked from overnight LB plates to inoculate 200 μL of LB containing corresponding antibiotic in a 2 mL 96-well block (Costar), and grown overnight at 1000 rpm in a VorTemp 56 bench top shaker (Labnet) at 37 °C. Then 4 μL of each overnight culture was added to 192 μL of LB containing corresponding antibiotics in a newly prepared 96-well block. Depending on the experiment, 4 μL of inducer was added to the newly prepared culture to reach the final concentrations of: 1, 3, 10, 30, 100, 300, 1000 μM of IPTG (Sigma-Aldrich) for Fig. 4B; 5, 10, 25, 50, 100, 200, 400 μM of cumate (Sigma-Aldrich) for Fig.4C; 1, 3, 10, 30, 100, 300, 1000 μM of 5-aminoimidazole-4-carboxamide ribonucleotide (Z, Sigma-Aldrich) for Fig. 4C; 1, 3, 10, 30, 100, 300, 1000 μM of 2-aminopurine (2-AP, Sigma-Aldrich) for Fig. 4D. The cells were then incubated at 1000 rpm in VorTemp at 37 °C for 8 hours before whole-cell fluorescence measurement.

For experiments using *S. oneidensis*, plasmids were transformed into electrocompetent cells. The cells were prepared by washing overnight-cultured cells 3 times with 10 % glycerol. Plasmids were electroporated into the cells using Gene Pulser Xcell electroporation system (Bio-Rad) with 1.2 kV, 25 μF, 200 Ω setting in 1 mm electroporation cuvettes (Fisher). *S. oneidensis* cells were recovered at 30 °C for 2 hours then plated on LB-agar plates containing 50 μg/mL kanamycin and incubated overnight at 30 °C.

For continuous culture experiments, single colonies were picked from the overnight-incubated LB-agar plates and inoculated in 50 mL glass culture tubes with 5 mL liquid LB containing corresponding antibiotics (LB-AB) and incubated in 37 °C shaking at 225 rpm. For every 24 hours, 5 μL of cell culture was inoculated to a new culture tube with 5 mL LB-AB (1:1000 dilution) till the end of the experiment.

### Whole-cell fluorescence measurement

Bulk fluorescence measurements were performed with 50 μL of culture diluted in 50 μL of phosphate-buffered saline (PBS, Fisher) in a 96-well plate using a microplate reader (Tecan Spark). The GFP was measured with 485/520 mm as excitation/emission wavelengths. The RFP was measured with 560/630 mm as excitation/emission wavelengths. Optical density at 600 nm (OD600) was also measured. Each 96-well block included two sets of controls; a media blank and *E. coli* transformed with empty control plasmids, or *S. oneidensis* without plasmids, referred to here as blank cells. Blank cells were used to determine autofluorescence levels. OD and fluorescence values for each colony were first corrected by subtracting the mean value of the media blank from the respective values of the experimental conditions. The ratio of the corrected fluorescence to the corrected OD (fluorescence/OD) was then calculated for each well. Autofluorescence was removed through the subtraction of fluorescence/OD of the blank cells.

For single-cell fluorescence analysis, single colonies were picked from the overnight-incubated LB-agar plates and inoculated in 50 mL glass culture tubes with 5 mL liquid LB containing 34 μg/mL chloramphenicol. After overnight culture in 37 °C, an aliquot was taken and diluted by PBS (1:100). The fluorescence distribution of the single cells in the sample were measured by flow cytometry (Sony SH800S cell sorter). Each measurement contains 100,000 event reads. The data was then processed and analyzed in the FlowJo software.

### Selection for functional RNA primer truncations

A DNA oligonucleotide library (Twist Bioscience) was ordered and then amplified through Polymerase Chain Reaction (PCR) using forward and reverse primer shown in Table S6, with the following conditions: 98 °C (30 s), 98 °C (15 s) 63 °C (45 s) 72 °C (15 s) for 13 cycles, 72 °C (5 min), 4 °C (forever). This amplified product was used to create the plasmid library containing different RNA primer truncations through Golden Gate assembly. The plasmid library was transformed into *E. coli* cells and then incubated at 37 °C overnight on LB-agar plates with antibiotic selection. Cells were removed from the plate and DNA plasmids were isolated using a DNA mini plasmid preparation (Qiagen). Using 10 ng of DNA plasmid as a template, the RNA primer (RNAII) region was amplified using primers also containing adapters for NGS sequencing using Amplicon-EZ (Genewiz). The following PCR conditions were used: 98 °C (30 s), 98 °C (15 s) 63 °C (45 s) 72 °C (15 s) for 20 cycles, 72 °C (5 min), 4 °C (forever). As a naïve control, the amplified PCR product used to construct the plasmid library before transformation was also prepared and sequenced in parallel. The relative enrichments were calculated by dividing the reads from each plasmid variant by the reads from the corresponding naïve control, which was then further normalized by the enrichment of the plasmid without truncation (0 nt).

### Growth rate characterization

Single colonies were picked from the overnight-incubated LB-agar plates and inoculated in 50 mL glass culture tubes with 5 mL liquid LB containing 34 μg/mL chloramphenicol (LB-AB). After overnight growth at 37 °C shaking at 225 rpm, an aliquot was taken and diluted 1:10000 with fresh liquid LB-AB and 200 μL of diluted sample was added to each well of a 96-well plate. The plate was then placed in the microplate reader (Tecan Spark) with the humidity cassette for continuous culture at 37°C shaking at 108 rpm. The OD600 of each sample was measured every 10 min for 20 hours.

### Plasmid loss characterization

Single colonies were picked from the overnight-incubated LB-agar plates and inoculated in 50 mL glass culture tubes with 5 mL liquid LB either containing (LB-AB) or lacking 34 μg/mL chloramphenicol (LB). An aliquot of the culture was then streaked on LB-agar plates without antibiotic present and grown overnight. For each biological replicate, 20 colonies were picked and transferred to a new LB-agar plate containing 34 μg/mL chloramphenicol and grown overnight. The survival rate of transferred colonies was then determined to characterize plasmid loss.

### Dynamic characterization through microfluidic device

The experiment was conducted in a microfluidic device with dial-a-wave (DAW) junction and hallway traps^43^. Microfluidic devices were constructed using standard techniques^44^. *E. coli* cells were transformed with IPTG-inducible pSynORI, or with both IPTG-inducible and cumate-inducible pSynORIs. Cells were grown in LB media overnight with the corresponding antibiotic. After overnight growth at 37 °C shaking at 225 rpm, an aliquot was taken and diluted 1:1000 into 25 mL fresh LB media with antibiotic. After 6 hours of growth when OD600 reached ∼0.4, 15 mL of culture were centrifuged at 2000 rcf for 8 minutes at 25 °C. The cell pellet was then resuspended in 10 mL prewarmed fresh LB media with antibiotics and 0.5% Tween 20 (Fisher) and loaded into the microfluidic device at 37 °C. 0.5% Tween 20 was used in all media to prevent cells from adhering to the channel without affecting the cell growth and fluorescence expression.

For cells with IPTG-inducible pSynORI, after cells were loaded, fresh LB media (20 mL in 60 mL syringe) with antibiotic and 0.5% Tween 20 were added to the channel with a flow velocity of 100 μm/s through the channel. After cells fulfilled the trap and stabilized, the channel’s input media was switched to fresh LB media (20 mL in 60 mL syringe) with 100 μM IPTG, 10 μL Oregon Green 488 dye (Fisher), corresponding antibiotic, and 0.5% Tween 20 with the same flow velocity. After 17 hours, the input media was switched back to LB media without IPTG till the end of the experiment.

For cells with both IPTG-inducible and cumate-inducible pSynORIs, after cells were loaded, fresh LB media (20 mL in 60 mL syringe) with 100 μM IPTG, antibiotics and 0.5% Tween 20 were added to the channel with a flow velocity of 50 μm/s through the channel. After cells fulfilled the trap and stabilized, the channel’s input media was switched to fresh LB media (20 mL in 60 mL syringe) with 200 μM cumate, corresponding antibiotics, and 0.5% Tween 20 with the same flow velocity. After 17.8 hours, the input media was switched back to LB media with 100 μM IPTG till the end of the experiment. Both types of media contained no dye, and the media switch time was recorded manually during the switch.

Phase contrast and fluorescence images were acquired under an inverted fluorescence microscope (Nikon) every 5 minutes at 100X magnification. For cells containing a single pSynORI the exposure times were set to 100ms for RFP fluorescence channel and 400ms for GFP fluorescence channel used for the dye. For cells containing a double pSynORI the exposure times were set to 100ms for both RFP and GFP fluorescence channels. Experimentally measured time course data was acquired and analyzed with ImageJ and MATLAB.

### Nanopore sequencing and data analysis

For sample preparation, single colonies were picked from the overnight-incubated LB-agar plates and inoculated in 50 mL glass culture tubes with 5 mL liquid LB containing corresponding antibiotics at 37 °C shaking at 225 rpm (LB-AB). For data shown in Fig. 4H, 100 μM IPTG and 200 μM cumate were also added to the liquid culture. Following overnight growth, DNA plasmids were isolated using a DNA mini plasmid preparation (Qiagen). The plasmid sample was then sent for nanopore sequencing (Plasmidsaurus).

Data was processed and displayed as follows. For quantification of multimer species in Fig. 2F and 3C, the mass ratio of multimers from the sequencing result was presented. Mass ratio indicates the distribution of nucleotides, as well as ORI copies, across the different multimer species. For multiplex plasmid reporting in Fig. 4H, the number of nanopore sequencing reads for each plasmid were directly acquired from Plasmidsaurus. For 6-plasmid characterization in Fig. 5C, the raw nanopore sequencing read from Plasmidsaurus was processed with alignment workflow (wf-alignment) on EPI2ME platform using all 6 plasmids as the reference. The alignment result is then used to calculate the percentage of plasmid composition.

## Data availability

All data are available in the main text or the supplementary materials.

## Acknowledgments

We would like to thank Dr. Peter Davenport for helpful discussions and insights regarding this work. We would like to thank Dr. Julius Lucks for providing pT181 DNA plasmids.

## Author contributions

BL, JC, and MRL conceived the project. Experiments were performed by BL and XP. BL and JC wrote the manuscript, and all authors provided feedback.

## Supplementary Materials

Figs. S1 to S13

Tables S1 to S7

Movies S1 to S2

Data S1

