## Supplementary material for "Engineering Plasmids with Synthetic Origins of Replication": Supp Information

### TABLE OF CONTENTS


| Item | Title | Page |
|---|---|---|
| Fig. S1 | Characterization of the promoters from pMB1 origin of replication | 3 |
| Fig. S2 | Schematic of the pT181 attenuator mechanism | 4 |
| Fig. S3 | Fluorescent characterization of pSynORI created using the pT181 transcriptional attenuator | 5 |
| Fig. S4 | Comparing the copy number of pSynORI in E. coli and S. oneidensis cells | 6 |
| Fig. S5 | Histograms of nanopore sequencing read lengths with or without cer site | 7 |
| Fig. S6 | Next-generation sequencing reads characterizing relative plasmid enrichment with truncated priming RNA library | 8 |
| Fig. S7 | Effect of truncated priming RNA on pSynORI copy number | 9 |
| Fig. S8 | Regulation of plasmid copy through the independent control of the target RNA and repressor sRNA via separate chemical inputs | 10 |
| Fig. S9 | Independent replication regulation of two orthogonal pSynORIs with inducible copy number control. | 11 |
| Fig. S10 | Characterization of the dynamics of pSynORIs with inducible copy number control | 12 |
| Fig. S11 | Investigation of pSynORI library's orthogonality and stability | 13 |
| Fig. S12 | Comparison of multi-plasmid co-transformation results between pSynORI and pMB1 | 14 |
| Fig. S13 | The schematics of representative plasmids used in this study | 15 |
| Table S1 | All plasmids used in this study | 16-21 |
| Table S2 | pSynORI plasmid sequence | 22-24 |
| Table S3 | The sequences of promoters used in this study | 25 |
| Table S4 | The sequences of riboswitches used in this study | 26 |
| Table S5 | The sequences of pT181 regulatory elements used in this study | 27-28 |
| Table S6 | Examples of DNA fragments used to create the plasmid library with priming RNA truncations | 29 |
| Table S7 | Golden Gate assembly primers used to replace the regulatory elements in pSynORI | 30 |

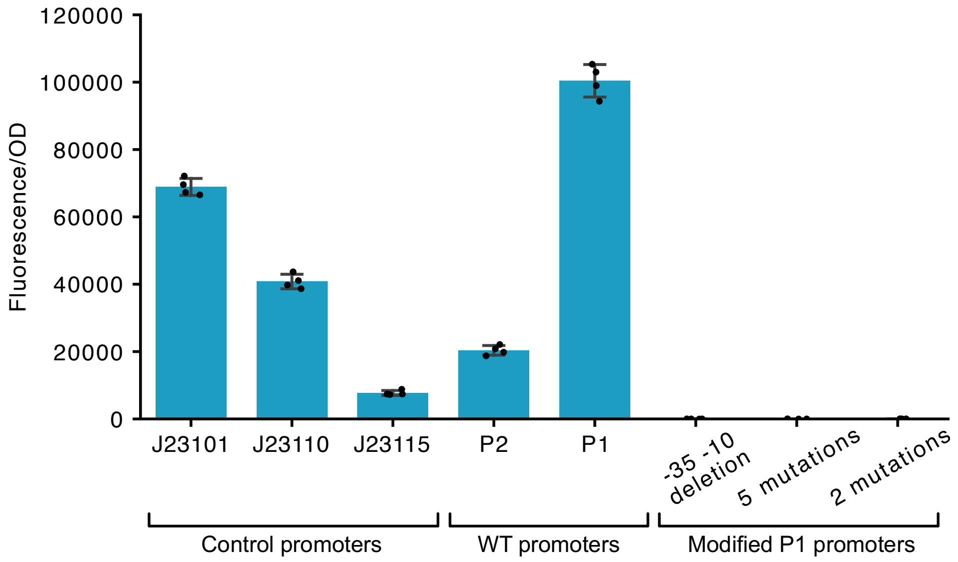


**Fig. S1. Characterization of the promoters from pMB1 origin of replication.** The strength of promoters was measured through fluorescence protein expression. Left to right: a set of synthetic control promoters; wild type promoters from pMB1 origin of replication; modified P1 promoters. Fluorescence characterization was performed (measured in units of fluorescence/optical density (OD) at 600 nm) in *E. coli* cells. Bars show mean and points show individual values of n = 4 biological replicates.

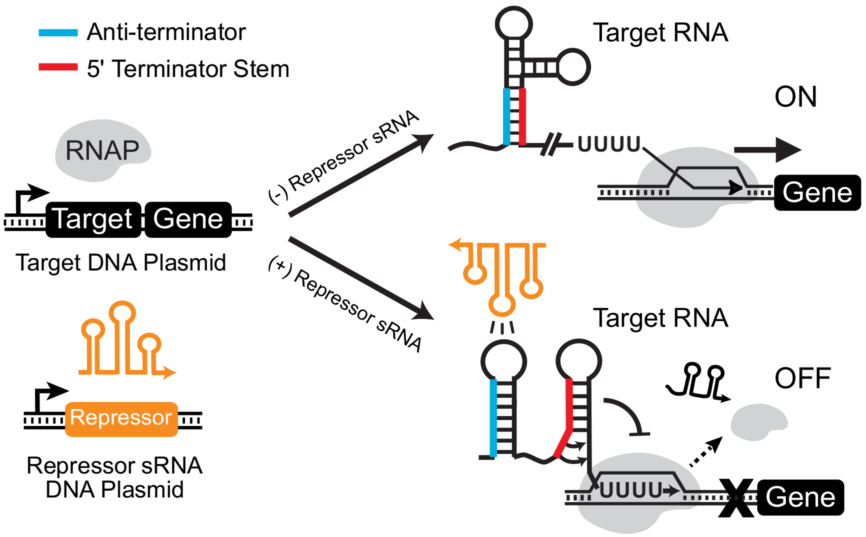


**Fig. S2. Schematic of the pT181 attenuator mechanism.** The pT181 attenuator consists of the target RNA and the repressor sRNA. The target RNA is inserted upstream of the gene to be controlled and contains within it a sequence that can form an intrinsic terminator hairpin. In the absence of repressor sRNA, the anti-terminator sequence sequesters the 5' terminator stem, preventing terminator formation and allowing downstream transcription (ON). When present, the repressor sRNA sequesters the anti-terminator of the target RNA, allowing terminator formation, and preventing downstream transcription (OFF).

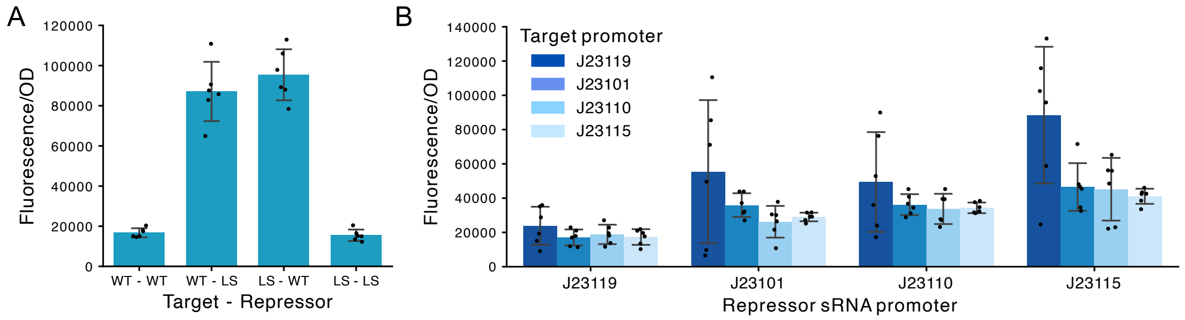


**Fig. S3. Fluorescent characterization of pSynORI created using the pT181 transcriptional attenuator.** The relative copy number characterization of the plasmids with (A) 4 combinations of pT181 target and repressor sRNA (WT and LS) and with (B) 16 combinations of pT181 target and repressor sRNA promoters (J23115, J23110, J23101, J23119) (cite). Relative copy number was characterized by encoding a constitutive RFP expression cassette onto the plasmid in *E. coli* cells (measured in units of Fluorescence/Optical density [OD]). Bars show mean and points show individual values of n = 6 biological replicates.

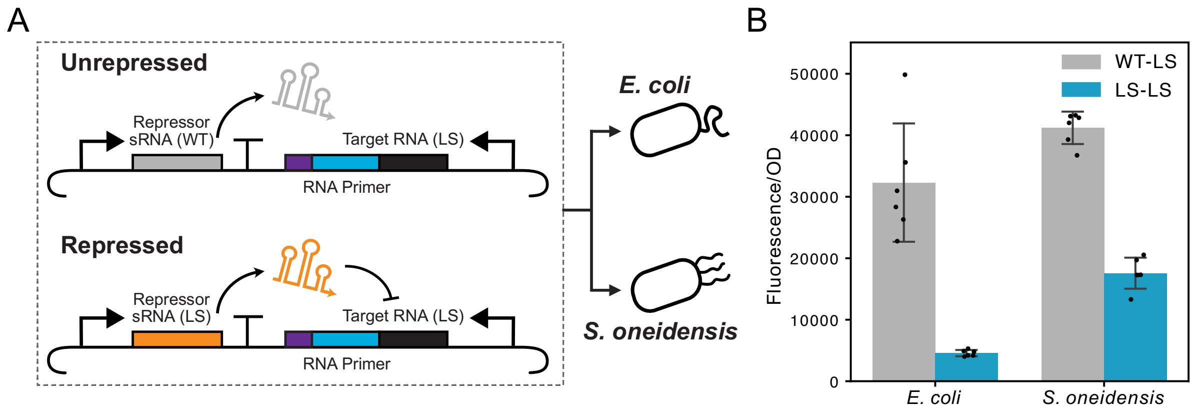


**Fig. S4. Comparing the copy number of pSynORI in *E. coli* and *S. oneidensis* cells.** (A) A pair of re-engineered pMB1 with matching and non-matching target RNA and repressor sRNA is transformed into *E. coli* and *S. oneidensis* cells. The LS variant of pT181 target has the interacting region modified so that it cannot be recognized by wild type (WT) repressor sRNA, leading to unrepressed plasmid replication. (B) Relative copy number was characterized by encoding a constitutive RFP expression cassette onto the plasmid in *E. coli* and *S. oneidensis* cells (measured in units of Fluorescence/Optical density [OD]). Bars show mean and points show individual values of n = 6 biological replicates.

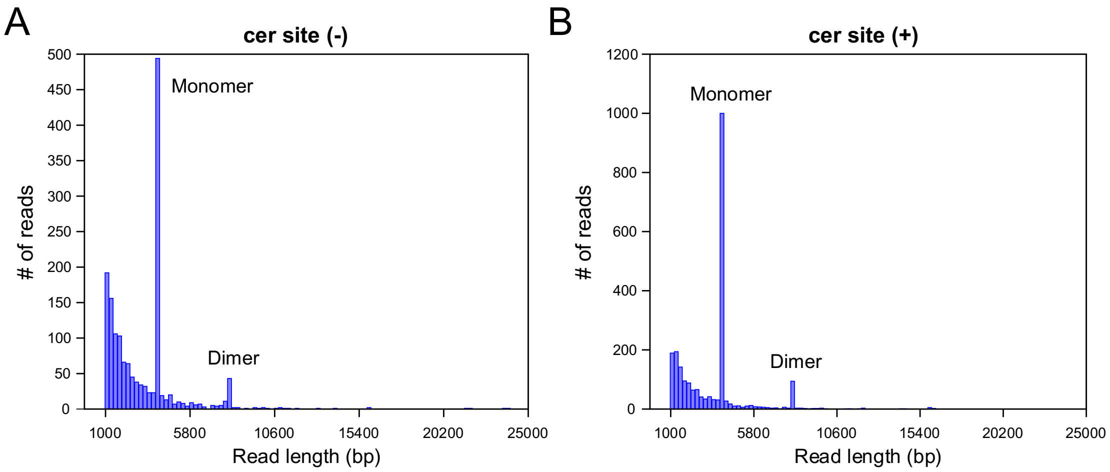


**Fig. S5. Histograms of nanopore sequencing read lengths with or without cer site.** The sequencing data from (A) the plasmid without cer site and (B) the plasmid with cer site. In both cases, the plasmid size is about 4000 base pairs (bp). The monomers are indicated by the first peak (~4000 bp). The dimers are indicated by the second peak (~8000bp). No significant decrease in dimer ratio was observed when cer site was added to the plasmid.

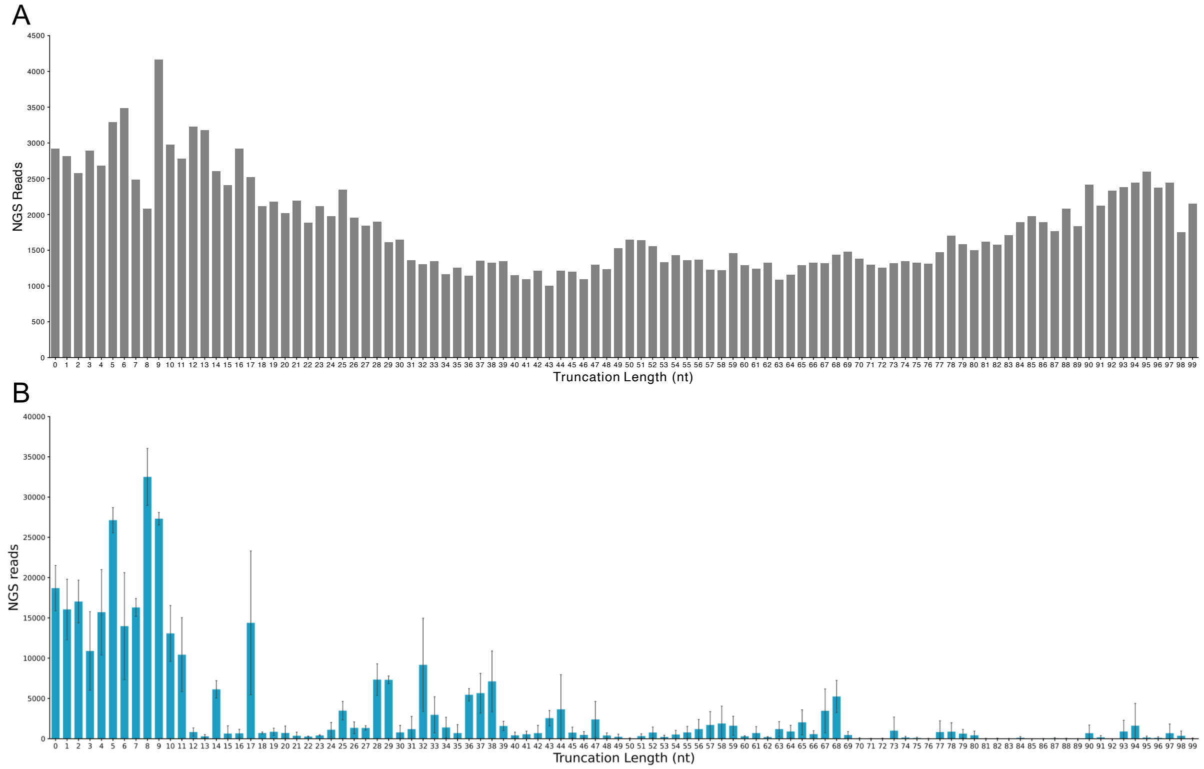


**Fig. S6. Next-generation sequencing reads characterizing relative plasmid enrichment with truncated priming RNA library.** (A) NGS reads of the initial DNA fragments used for plasmid cloning, serving as a control to calculate relative plasmid enrichment. (B) NGS reads characterizing the abundance of each plasmid design with priming RNA truncation after transformation, selection, and plasmid isolation. Bars show mean and the error bars show the standard deviation of n = 3 biological replicates.

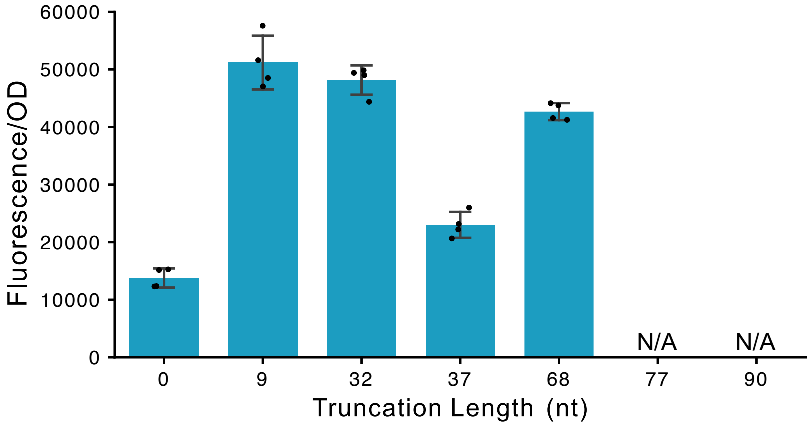


**Fig. S7. Effect of truncated priming RNA on pSynORI copy number.** The relative copy number characterization of plasmids with specific truncation length of priming RNA. 68-nt truncation is located at the junction between stem loop II and III. 77-nt and 90-nt truncation are inside the stem loop III and could not be cloned and is labeled as not available (N/A). Relative copy number was characterized by encoding a constitutive RFP expression cassette onto the plasmid in *E. coli* cells (measured in units of Fluorescence/Optical density [OD]). Bars show mean and points show individual values of n = 4 biological replicates.

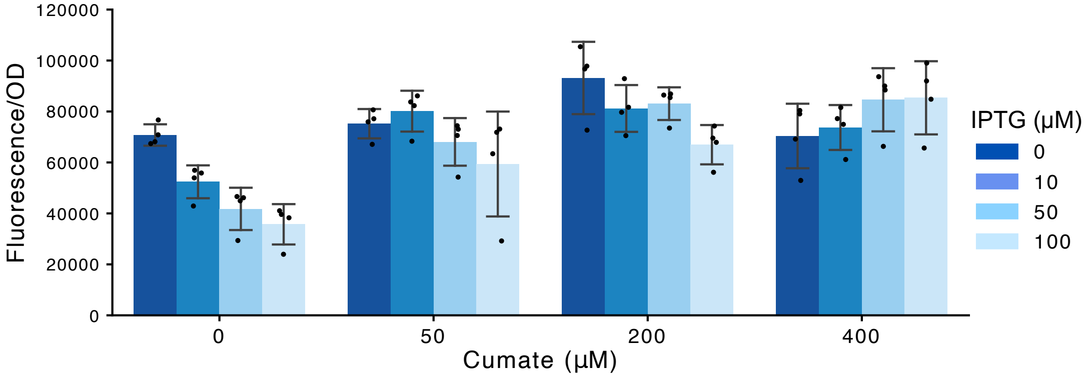


**Fig. S8. Regulation of plasmid copy through the independent control of the target RNA and repressor sRNA via separate chemical inputs.** The plasmid copy is up-regulated by cumate through target RNA transcription and down-regulated by IPTG through repressor sRNA transcription. Relative copy number was characterized by encoding a constitutive RFP expression cassette onto the plasmid in *E. coli* cells (measured in units of Fluorescence/Optical density [OD]). Bars show mean and points show individual values of n = 4 biological replicates.

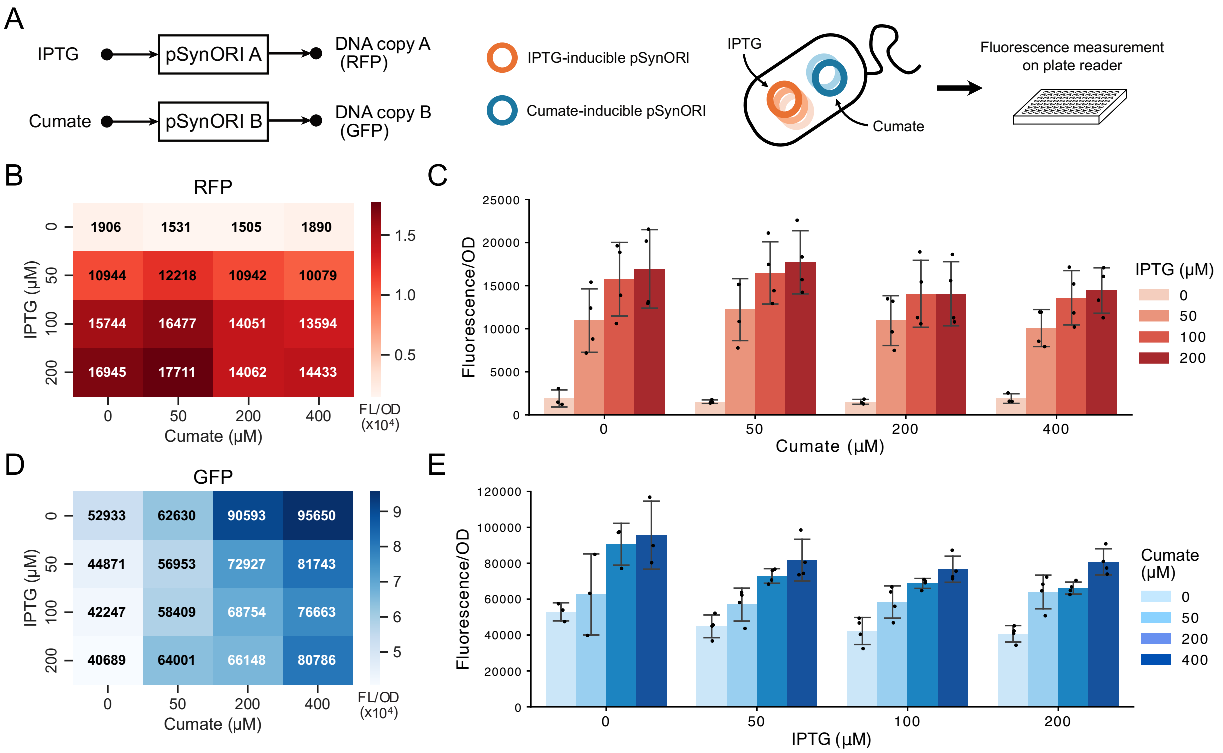


**Fig. S9. Independent replication regulation of two orthogonal pSynORIs with inducible copy number control.** (A) A pair of inducible pSynORI plasmids carrying GFP and RFP co-transformed into *E. coli* cell. (B) Matrix and (C) Bar plots showing the relative copy number characterization of IPTG-inducible plasmids as RFP across different cumate and IPTG concentration combinations. Bars show mean and points show individual values of n = 4 biological replicates. (D) Matrix and (E) Bar plots showing relative copy number of cumate-inducible plasmids expressing GFP across different cumate and IPTG concentration combinations. Bars show mean and points show individual values of n = 4 biological replicates.

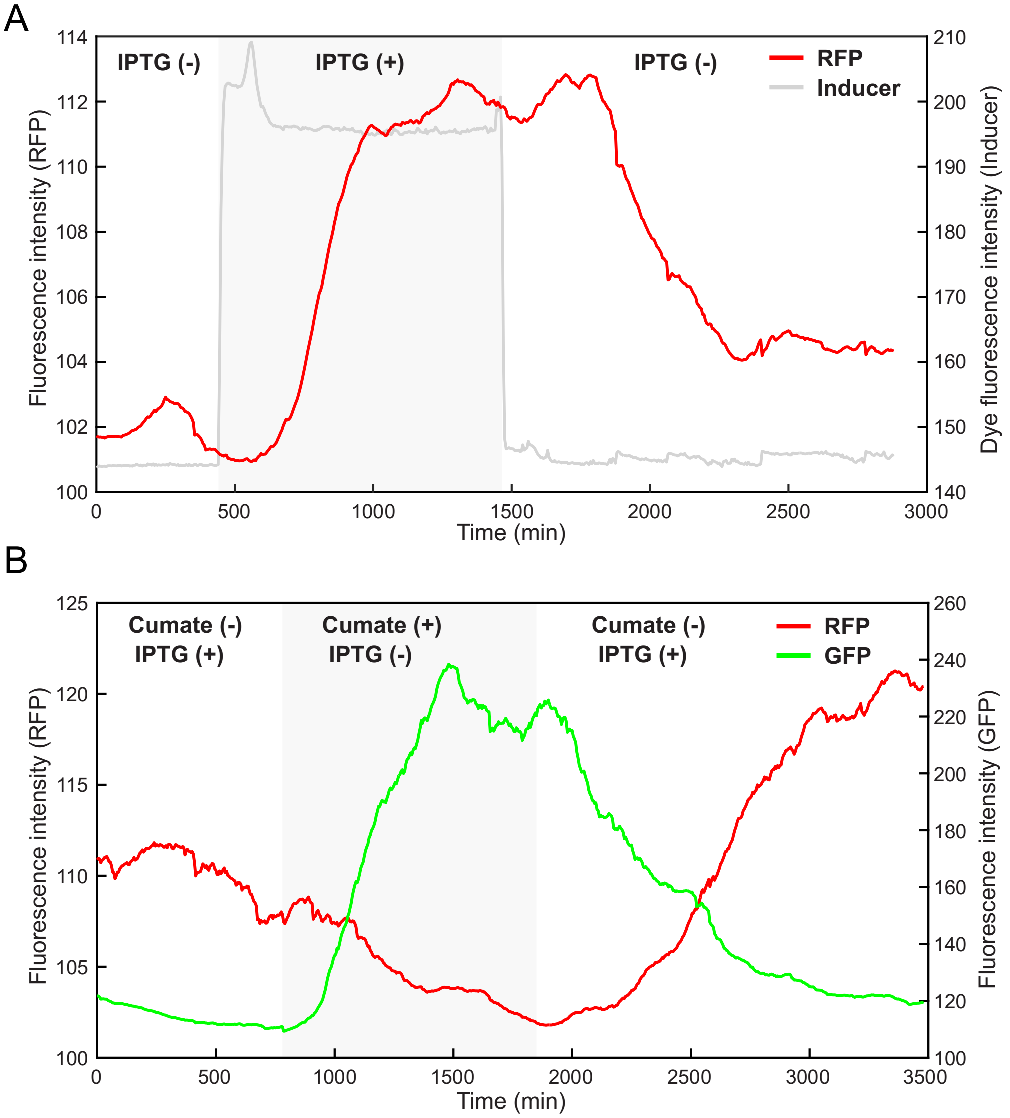


**Fig. S10. Characterization of the dynamics of pSynORIs with inducible copy number control.** (A) The IPTG-inducible pSynORI carrying constitutive expressed RFP was transformed into *E. coli* cells. The cells were put into a microfluid device, and fluorescence was measured using a microscope for 48 hours. The average fluorescence intensity in the cell trap was calculated and shown as a red line. The induction condition was indicated by a fluorescent dye added to IPTG (+) media, shown as a grey line. 100 µM IPTG was added at 7.5 hours and removed at the 24.5 hours. The time of the induction window is indicated by the grey shade. (B) The IPTG-inducible pSynORI carrying RFP and the cumate-inducible pSynORI carrying GFP were transformed into *E. coli* cells. The cells were put into a microfluid device, and fluorescence was measured using a microscope for 58 hours. The average RFP and GFP fluorescence intensity in the cell trap was calculated and shown as a red line and green line separately. Induction condition with either 100 µM IPTG or 200 µM cumate was switched at 13 hours and 30.8 hours, which is indicated by the grey shade.

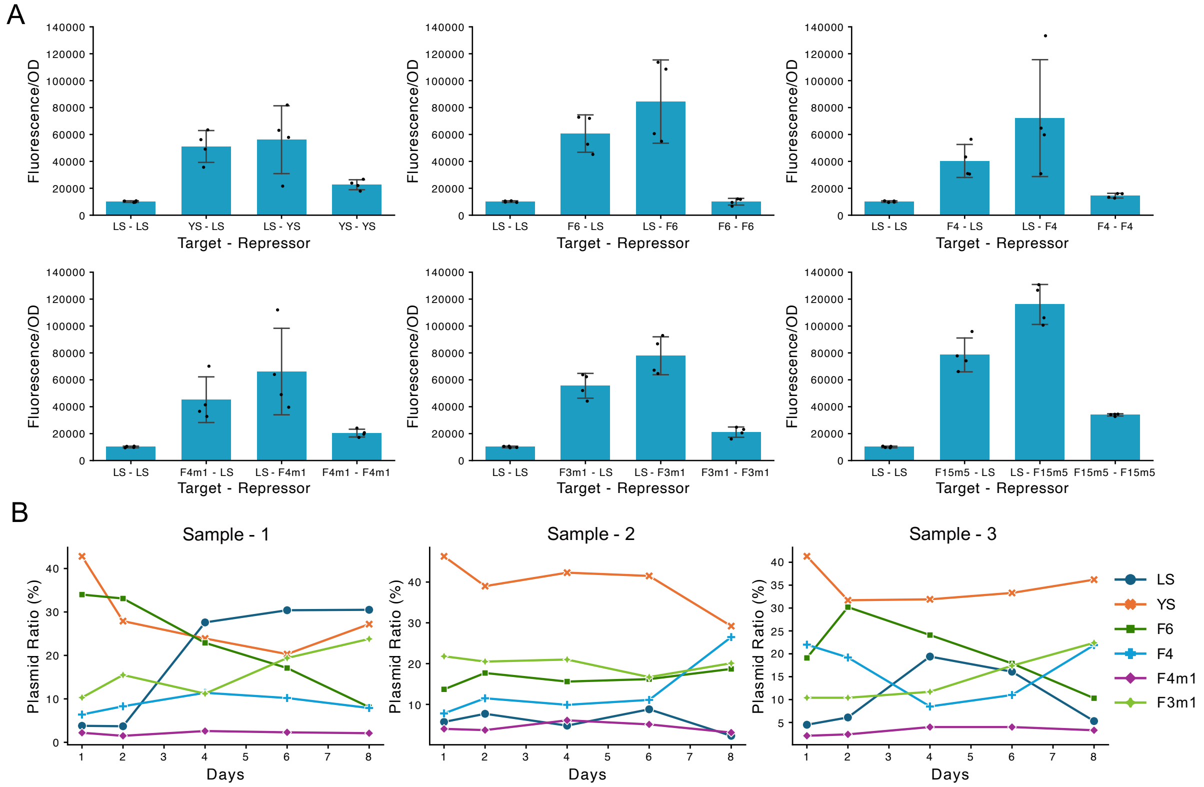


**Fig. S11. Investigation of pSynORI library's orthogonality and stability.** (A) The relative copy number characterization of the plasmids with combinations of pT181 target and repressor sRNA (LS, YS, F6, F4, F4m1, F3m1 to LS). Bars show mean and points show individual values of n = 4 biological replicates. (B) The composition ratio of 6 pSynORI plasmids in 3 biological replicate samples during an 8-day continuous culture.

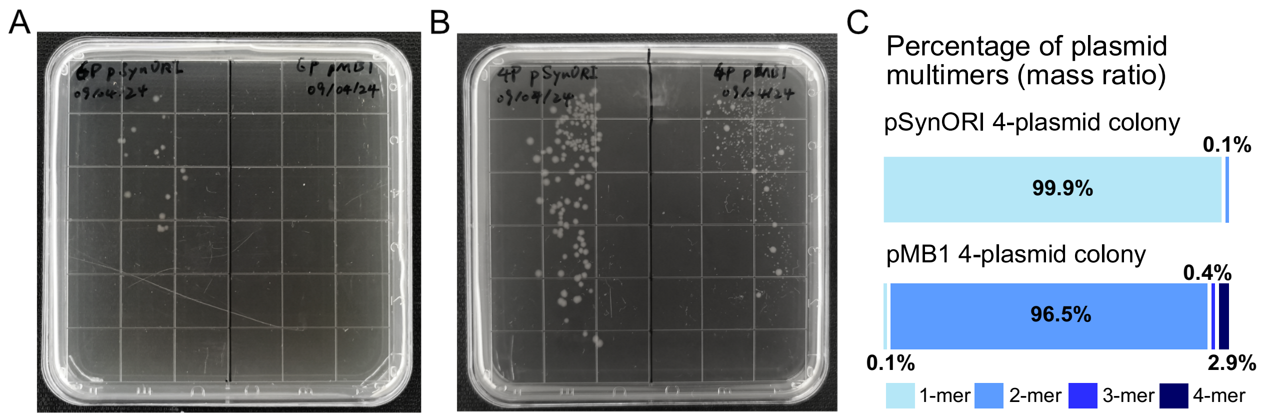


**Fig. S12. Comparison of multi-plasmid co-transformation results between pSynORI and pMB1.** (A) 6 pSynORI (LS, YS, F6, F4) (left side of plate) and 6 pMB1 control plasmids (right side of plate) or (B) 4 pSynORI (LS, YS, F6, F4, F4m1, F3m1) (left side of plate) and 4 pMB1 control plasmids (right side of plate) were co-transformed into chemical competent *E. coli* (NEB Turbo) and streaked on the LB-agar plate with corresponding antibiotics. Equal weight of plasmids (400 ng of each) and the same protocol was used to achieve comparable transformation efficiency. Left half of each plate is the result of pSynORI plasmid co-transformation. Right half of each plate is the result of pMB1 control plasmid transformation. (C) From the pMB1 4 plasmid transformation, 3 out of 5 picked colonies, all large, grew in liquid culture. Sequencing one of these cultures revealed a high rate of dimer formation, despite all plasmids being confirmed as monomers before transformation.

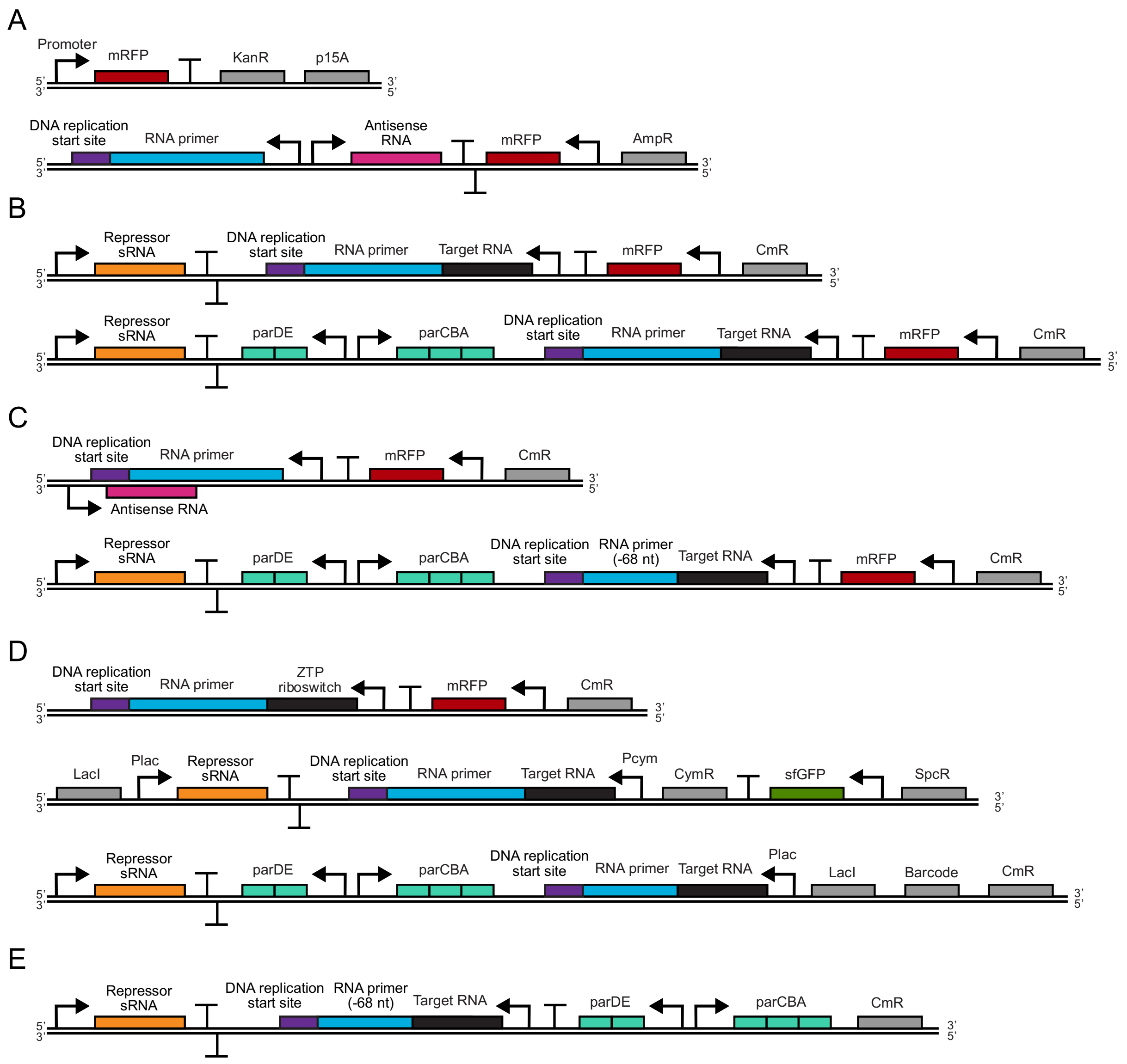


**Fig. S13. The schematics of representative plasmids used in this study.** (A-E) Key plasmids used in Figure 1-5 correspondingly.

**Table S1.**
**All plasmids used in this study.** CmR: chloramphenicol resistance gene, AmpR: ampicillin resistance gene, SpcR: spectinomycin resistance gene, KanR: kanamycin resistance gene, TetR: tetracycline resistance gene, AprR: apramycin resistance gene, pMB1: origin of replication, p15A: origin of replication, pSC101: origin of replication, SynORI: synthetic origin of replication, RBS: ribosome binding site, J23119, J23101, J23110, J23115: constitutive promoters from Anderson promoter collection.

| Plasmid | Description | Plasmid Architect | Figure |
|---|---|---|---|
| JEC1521 | P2(WT)-RFP | P2(WT)-RBS-RFP-Terminator-KanR-p15A | Fig1C, figS1 |
| JEC1522 | P2(2-mutation)-RFP | P2(2-mutation)-RBS-RFP-Terminator-KanR-p15A | Fig1C, figS1 |
| JEC1523 | P2(5-mutation)-RFP | P2(5-mutation)-RBS-RFP-Terminator-KanR-p15A | Fig1C, figS1 |
| JEC1524 | P2(-10 -35 deletion)-RFP | P2(-10 -35 deletion)-RBS-RFP-Terminator-KanR-p15A | Fig1C, figS1 |
| JEC1525 | P1(WT)-RFP | P1(WT)-RBS-RFP-Terminator-KanR-p15A | figS1 |
| JEC1526 | J23101-RFP | J23101-RBS-RFP-Terminator-KanR-p15A | figS1 |
| JEC1527 | J23110-RFP | J23110-RBS-RFP-Terminator-KanR-p15A | figS1 |
| JEC1528 | J23115-RFP | J23115-RBS-RFP-Terminator-KanR-p15A | figS1 |
| JEC1529 | pMB1(WT) | pMB1-AmpR-RFP | Fig1C |
| JEC1530 | Refactored pMB1 (2-mutation) | Inhibiting RNA-P2-P1-RNA Primer (2-mutation P2)-RFP-AmpR | Fig1C |
| JEC1531 | Refactored pMB1 (5-mutation) | Inhibiting RNA-P2-P1-RNA Primer (5-mutation P2)-RFP-AmpR | Fig1C |
| JEC1532 | Refactored pMB1 (-10 -35 deletion) | Inhibiting RNA-P2-P1-RNA Primer (-10 -35 deletion P2)-RFP-AmpR | Fig1C |
| JEC1533 | Reengineered pMB1 (Repressor - WT/Target-WT) | Repressor (WT)-J23119-J23110-Target (WT)-RNA Primer-RFP-CmR | Fig2B, figS3A |
| JEC1534 | Reengineered pMB1 (Repressor - LS/Target-WT) | Repressor (LS)-J23119-J23110-Target (WT)-RNA Primer-RFP-CmR | Fig2B, figS3A |
| JEC1535 | Reengineered pMB1 (Repressor - WT/Target-LS) | Repressor (WT)-J23119-J23110-Target (LS)-RNA Primer-RFP-CmR | Fig2B, figS3A |
| JEC1536 | Reengineered pMB1 (Repressor - LS/Target-LS) | Repressor (LS)-J23119-J23110-Target (LS)-RNA Primer-RFP-CmR | Fig2B, figS3A |

| | | | |
|---|---|---|---|
| JEC1537 | Reengineered pMB1 (Repressor-J23119/Target-J23119) | Repressor (WT)-J23119-J23119-Target (WT)-RNA Primer-RFP-AmpR | Fig2C, figS3B |
| JEC1538 | Reengineered pMB1 (Repressor-J23119/Target-J23101) | Repressor (WT)-J23119-J23101- Target (WT)-RNA Primer-RFP-AmpR | Fig2C, figS3B |
| JEC1539 | Reengineered pMB1 (Repressor-J23119/Target-J23110) | Repressor (WT)-J23119-J23110- Target (WT)-RNA Primer-RFP-AmpR | Fig2C, figS3B |
| JEC1540 | Reengineered pMB1 (Repressor-J23119/Target-J23115) | Repressor (WT)-J23119-J23115- Target (WT)-RNA Primer-RFP-AmpR | Fig2C, figS3B |
| JEC1541 | Reengineered pMB1 (Repressor-J23101/Target-J23119) | Repressor (WT)-J23101-J23119- Target (WT)-RNA Primer-RFP-AmpR | Fig2C, figS3B |
| JEC1542 | Reengineered pMB1 (Repressor-J23101/Target-J23101) | Repressor (WT)-J23101-J23101- Target (WT)-RNA Primer-RFP-AmpR | Fig2C, figS3B |
| JEC1543 | Reengineered pMB1 (Repressor-J23101/Target-J23110) | Repressor (WT)-J23101-J23110- Target (WT)-RNA Primer-RFP-AmpR | Fig2C, figS3B |
| JEC1544 | Reengineered pMB1 (Repressor-J23101/Target-J23115) | Repressor (WT)-J23101-J23115- Target (WT)-RNA Primer-RFP-AmpR | Fig2C, figS3B |
| JEC1545 | Reengineered pMB1 (Repressor-J23110/Target-J23119) | Repressor (WT)-J23110-J23119- Target (WT)-RNA Primer-RFP-AmpR | Fig2C, figS3B |
| JEC1546 | Reengineered pMB1 (Repressor-J23110/Target-J23101) | Repressor (WT)-J23110-J23101- Target (WT)-RNA Primer-RFP-AmpR | Fig2C, figS3B |
| JEC1547 | Reengineered pMB1 (Repressor-J23110/Target-J23110) | Repressor (WT)-J23110-J23110- Target (WT)-RNA Primer-RFP-AmpR | Fig2C, figS3B |

| | | | |
|---|---|---|---|
| JEC1548 | Reengineered pMB1 (Repressor-J23110/Target-J23115) | Repressor (WT)-J23110-J23115- Target (WT)-RNA Primer-RFP-AmpR | Fig2C, figS3B |
| JEC1549 | Reengineered pMB1 (Repressor-J23115/Target-J23119) | Repressor (WT)-J23115-J23119- Target (WT)-RNA Primer-RFP-AmpR | Fig2C, figS3B |
| JEC1550 | Reengineered pMB1 (Repressor-J23115/Target-J23101) | Repressor (WT)-J23115-J23101- Target (WT)-RNA Primer-RFP-AmpR | Fig2C, figS3B |
| JEC1551 | Reengineered pMB1 (Repressor-J23115/Target-J23110) | Repressor (WT)-J23115-J23110- Target (WT)-RNA Primer-RFP-AmpR | Fig2C, figS3B |
| JEC1552 | Reengineered pMB1 (Repressor-J23115/Target-J23115) | Repressor (WT)-J23115-J23115- Target (WT)-RNA Primer-RFP-AmpR | Fig2C, figS3B |
| JEC1553 | Reengineered pMB1 (Divergent) | Repressor (WT)-J23119-J23110-Target (WT)-RNA Primer-RFP-CmR | Fig2D |
| JEC1554 | Reengineered pMB1 (Tandem down) | J23119-Repressor (WT)-J23110-Target (WT)-RNA Primer-RFP-CmR | Fig2D |
| JEC1555 | Reengineered pMB1 (Tandem up) | Repressor (WT)-J23119-RNA Primer-Target (WT)-J23110-RFP-CmR | Fig2D |
| JEC1556 | Reengineered pMB1 (Convergent) | J23119-Repressor (WT)-RNA Primer-Target (WT)-J23110-RFP-CmR | Fig2D |
| JEC1557 | Reengineered pMB1 (Unrepressed) | J23119-Repressor (WT)-RNA Primer-Target (LS)-J23110-GFP-SpcR | figS4 |
| JEC1558 | Reengineered pMB1 (Repressed) | J23119-Repressor (LS)-RNA Primer-Target (LS)-J23110-GFP-SpcR | figS4, Fig5A |
| JEC1559 | Reengineered pMB1 (parCBA-) | J23119-Repressor (LS)-RNA Primer-Target (LS)-J23110-RFP-CmR | Fig2F |
| JEC1560 | Reengineered pMB1 (parCBA+) | J23119-Repressor (LS)-parDE-parCBA-RNA Primer-Target (LS)-J23110-RFP-CmR | Fig2F |
| JEC1561 | Reengineered pMB1 (cer-) | J23119-Repressor (LS)-RNA Primer-Target (LS)-J23101-RFP-CmR | figS5A |
| JEC1562 | Reengineered pMB1 (cer+) | J23119-Repressor (LS)-cer-RNA Primer-Target (LS)-J23101-RFP-CmR | figS5B |

| | | | |
|---|---|---|---|
| JEC1563 | Reengineered pMB1 (Full length) | J23119-Repressor (WT)-RNA Primer (Full length)-Target (WT)-J23110-RFP-CmR | figS7 |
| JEC1564 | Reengineered pMB1 (-9 nt) | J23119-Repressor (WT)-RNA Primer (-9 nt)-Target (WT)-J23110-RFP-CmR | figS7 |
| JEC1565 | Reengineered pMB1 (-32 nt) | J23119-Repressor (WT)-RNA Primer (-32 nt)-Target (WT)-J23110-RFP-CmR | figS7 |
| JEC1566 | Reengineered pMB1 (-37 nt) | J23119-Repressor (WT)-RNA Primer (-37 nt)-Target (WT)-J23110-RFP-CmR | figS7 |
| JEC1567 | Reengineered pMB1 (-68 nt) | J23119-Repressor (WT)-RNA Primer (-68 nt)-Target (WT)-J23110-RFP-CmR | figS7 |
| JEC1568 | pMB1 control | pMB1-RFP-CmR | Fig3A-E |
| JEC1569 | pSynORI | J23119-Repressor (LS)-parDE-parCBA-RNA Primer (-68nt)-Target (LS)-J23110-RFP-CmR | Fig3A-E |
| JEC1570 | Reengineered pMB1 (IPTG-inducible) | J23119-Repressor (LS)-RNA Primer-Target (LS)-Plac-LacI-RFP-CmR | Fig4B, figS9B-E, figS10A-B |
| JEC1571 | Reengineered pMB1 (Cumate-inducible) | J23119-Repressor (LS)-RNA Primer-Target (LS)-Pcym-CymR-GFP-SpcR | Fig4C, figS9B-E, figS10B |
| JEC1572 | Reengineered pMB1 (Z-inducible) | RNA Primer-ZTP riboswitch-J23110-RFP-CmR | Fig4D |
| JEC1573 | Reengineered pMB1 (2AP-inducible) | RNA Primer-2AP riboswitch-J23110-RFP-CmR | Fig4E |
| JEC1574 | Reengineered pMB1 (Dual-inducible) | LacI-Plac-Repressor (WT)-RNA Primer-Target (WT)-Pcym-CymR-GFP-SpcR | Fig4F, figS8 |
| JEC1575 | pSynORI (IPTG-inducible) | J23119-Repressor (LS)-parDE-parCBA-RNA Primer-Target (LS)-Plac-LacI-Barcode1-CmR | Fig4H |
| JEC1576 | pSynORI (Cumate-inducible) | J23119-Repressor (LS)-parDE-parCBA-RNA Primer-Target (LS)-Pcym-CymR-Barcode2-SpcR | Fig4H |
| JEC1577 | pSC101 control | Barcode3-pSC101-KanR | Fig4H |
| JEC1578 | Reengineered pMB1 (Repressor-LS/Target-YS) | J23119-Repressor (LS)-RNA Primer-Target (YS)-J23110-GFP-SpcR | Fig5A, figS11A |
| JEC1579 | Reengineered pMB1 (Repressor-LS/Target-F6) | J23119-Repressor (LS)-RNA Primer-Target (F6)-J23110-GFP-SpcR | Fig5A, figS11A |
| JEC1580 | Reengineered pMB1 (Repressor-LS/Target-F4m1) | J23119-Repressor (LS)-RNA Primer-Target (F4m1)-J23110-GFP-SpcR | Fig5A, figS11A |

| | | | |
|---|---|---|---|
| JEC1581 | Reengineered pMB1 (Repressor-LS/Target-F4) | J23119-Repressor (LS)-RNA Primer-Target (F4)-J23110-GFP-SpcR | Fig5A, figS11A |
| JEC1582 | Reengineered pMB1 (Repressor-LS/Target-F3m1) | J23119-Repressor (LS)-RNA Primer-Target (F3m1)-J23110-GFP-SpcR | Fig5A, figS11A |
| JEC1583 | Reengineered pMB1 (Repressor-LS/Target-F15m5) | J23119-Repressor (LS)-RNA Primer-Target (F15m5)-J23110-GFP-SpcR | Fig5A, figS11A |
| JEC1584 | Reengineered pMB1 (Repressor-YS/Target-LS) | J23119-Repressor (YS)-RNA Primer-Target (LS)-J23110-GFP-SpcR | Fig5A, figS11A |
| JEC1585 | Reengineered pMB1 (Repressor-F6/Target-LS) | J23119-Repressor (F6)-RNA Primer-Target (LS)-J23110-GFP-SpcR | Fig5A, figS11A |
| JEC1586 | Reengineered pMB1 (Repressor-F4m1/Target-LS) | J23119-Repressor (F4m1)-RNA Primer-Target (LS)-J23110-GFP-SpcR | Fig5A, figS11A |
| JEC1587 | Reengineered pMB1 (Repressor-F4/Target-LS) | J23119-Repressor (F4)-RNA Primer-Target (LS)-J23110-GFP-SpcR | Fig5A, figS11A |
| JEC1588 | Reengineered pMB1 (Repressor-F3m1/Target-LS) | J23119-Repressor (F3m1)-RNA Primer-Target (LS)-J23110-GFP-SpcR | Fig5A, figS11A |
| JEC1589 | Reengineered pMB1 (Repressor-F15m5/Target-LS) | J23119-Repressor (F15m5)-RNA Primer-Target (LS)-J23110-GFP-SpcR | Fig5A, figS11A |
| JEC1590 | Reengineered pMB1 (Repressor-YS/Target-YS) | J23119-Repressor (YS)-RNA Primer-Target (YS)-J23110-GFP-SpcR | Fig5A, figS11A |
| JEC1591 | Reengineered pMB1 (Repressor-F6/Target-F6) | J23119-Repressor (F6)-RNA Primer-Target (F6)-J23110-GFP-SpcR | Fig5A, figS11A |
| JEC1592 | Reengineered pMB1 (Repressor-F4m1/Target-F4m1) | J23119-Repressor (F4m1)-RNA Primer-Target (F4m1)-J23110-GFP-SpcR | Fig5A, figS11A |
| JEC1593 | Reengineered pMB1 (Repressor-F4/Target-F4) | J23119-Repressor (F4)-RNA Primer-Target (F4)-J23110-GFP-SpcR | Fig5A, figS11A |
| JEC1594 | Reengineered pMB1 (Repressor-F3m1/Target-F3m1) | J23119-Repressor (F3m1)-RNA Primer-Target (F3m1)-J23110-GFP-SpcR | Fig5A, figS11A |

| | | | |
|---|---|---|---|
| JEC1595 | Reengineered pMB1 (Repressor-F15m5/Target-F15m5) | J23119-Repressor (F15m5)-RNA Primer-Target (F15m5)-J23110-GFP-SpcR | Fig5A, figS11A |
| JEC1596 | pSynORI (LS) | J23119-Repressor (LS)-RNA Primer (-68nt)-Target (LS)-J23110-parDE-parCBA-CmR | Fig5C, figS11B, figS12A-C |
| JEC1597 | pSynORI (YS) | J23119-Repressor (YS)-RNA Primer (-68nt)-Target (YS)-J23110-parDE-parCBA-SpcR | Fig5C, figS11B, figS12A-C |
| JEC1598 | pSynORI (F6) | J23119-Repressor (F6)-RNA Primer (-68nt)-Target (F6)-J23110-parDE-parCBA-KanR | Fig5C, figS11B, figS12A-C |
| JEC1599 | pSynORI (F4) | J23119-Repressor (F4)-RNA Primer (-68nt)-Target (F4)-J23110-parDE-parCBA-AmpR | Fig5C, figS11B, figS12A-C |
| JEC1600 | pSynORI (F4m1) | J23119-Repressor (F4m1)-RNA Primer (-68nt)-Target (F4m1)-J23110-parDE-parCBA-TetR | Fig5C, figS11B, figS12A |
| JEC1601 | pSynORI (F3m1) | J23119-Repressor (F3m1)-RNA Primer (-68nt)-Target (F3m1)-J23110-parDE-parCBA-AprR | Fig5C, figS11B, figS12A |
| JEC1602 | pMB1 CmR control | pMB1-CmR | figS12A-C |
| JEC1603 | pMB1 SpcR control | pMB1-SpcR | figS12A-C |
| JEC1604 | pMB1 KanR control | pMB1-KanR | figS12A-C |
| JEC1605 | pMB1 AmpR control | pMB1-AmpR | figS12A-C |
| JEC1606 | pMB1 TetR control | pMB1-TetR | figS12B |
| JEC1607 | pMB1 AprR control | pMB1-AprR | figS12B |

**Table S2.**
**pSynORI plasmid sequence.**

| Name | DNA sequence |
|---|---|
| pSynORI<br>(pBL724) CmR-<br>J23119-<br>Repressor (LS)-<br>Terminator-<br>parDE-parCBA-<br>RNA Primer-<br>Target (LS)-<br>J23110-RFP | AATAAAAAACGCCCGGCGGCAACCGAGCGTTCTGAACAAATCCAGATGGAGTTCT<br>GAGGTCATTACTGGATCTATCAACAGGAGTCCAAGCGAGCTCGATATCAAATTAC<br>GCCCCGCCCTGCCACTCATCGCAGTACTGTTGTAATTCATTAAGCATTCTGCCGAC<br>ATGGAAGCCATCACAAACGGCATGATGAACCTGAATCGCCAGCGGCATCAGCAC<br>CTTGTCGCCTTGCGTATAATATTTGCCCATGGTGAAAACGGGGGCGAAGAAGTTG<br>TCCATATTGGCCACGTTTAAATCAAAACTGGTGAAACTCACCCAGGGATTGGCTG<br>AGACGAAAAACATATTCTCAATAAACCCTTTAGGGAAATAGGCCAGGTTTTCACC<br>GTAACACGCCACATCTTGCGAATATATGTGTAGAAACTGCCGGAAATCGTCGTGG<br>TATTCACTCCAGAGCGATGAAAACGTTTCAGTTTGCTCATGGAAAACGGTGTAAC<br>AAGGGTGAACACTATCCCATATCACCAGCTCACCGTCTTTCATTGCCATACGAAAT<br>TCCGGATGAGCATTCATCAGGCGGGCAAGAATGTGAATAAAGGCCGGATAAAAC<br>TTGTGCTTATTTTTCTTTACGGTCTTTAAAAAGGCCGTAATATCCAGCTGAACGGT<br>CTGGTTATAGGTACATTGAGCAACTGACTGAAATGCCTCAAAATGTTCTTTACGAT<br>GCCATTGGGATATATCAACGGTGGTATATCCAGTGATTTTTTTCTCCATTTTAGCTT<br>CCTTAGCTCCTGAAAATCTCGATAACTCAAAAAATACGCCCGGTAGTGATCTTATT<br>TCATTATGGTGAAAGTTGGAACCTCTTACGTGCCCTGTCAGACCAAGTTTACTCAT<br>ATATACTTTAGATTGATTTAAAACTTCATTTTTAATTTAAAAGGATCTAGGTGAAG<br>ATCCTTTTTGATAATCTCATGACCAAAATCCCTTAACGTGAGTTTTCGTTCCACTG<br>AGCGTCAGACCCCGTAGAAAAGATCAAAGGATCTTCACGGAGGAATGGGTAGGT<br>ATTGAGTCTTCTTGACAGCTAGCTCAGTCCTAGGTATAATACTAGTCGACATACAA<br>GATTATAAAAACAACTCAGTGTTTTTTTCTTTGAATGATGTCGTTCTGCAACTTTG<br>GCGAGGGACAGAGCGACTCCTTTTTATTTGGATCTGAAGCTTGGGCCCGAACAAA<br>AACTCATCTCAGAAGAGGATCTGAATAGCGCCGTCGACCATCATCATCATCATCA<br>TTGAGTTTAAACGGTCTCCAGCTTGGCTGTTTTGGCGGATGAGAGAAGATTTTCAG<br>CCTGATACAGATTAAATCAGAACGCAGAAGCGGTCTGATAAAACAGAATTTGCCT<br>GGCGGCAGTAGCGCGGTGGTCCCACCTGACCCCATGCCGAACTCAGAAGTGAAAC<br>GCCGTAGCGCCGATGGTAGTGTGGGGTCTCCCCATGCGAGAGTAGGGAACTGCCA<br>GGCATCAAATAAAACGAAAGGCTCAGTCGAAAGACTGGGCCTTTCGTTTTATCTG<br>TTGTTTGTCGGTGAACTCGCACTTAACATCAATCTAATTATATATCATTATTACCGT<br>ACGCCATCAGGACGTTGTGAGTGGCGCGATTTTTAGCGGCTGAAATCAGCCCTTG<br>AGCCTGTCGGCAAGTCGCGTCATGAGGTCCATGCGCTCATGCAGGATCGCCACGA<br>CCAACGCGGGTTCGCCCGCACGCGGCAGGCAAAAAACGTAGTGGTGTTCGCAGC<br>GGGCCATCCGCAGCGCGGGAAAGAGTTCGCTCATGTCCTTAAACGGGCCTTCGCC<br>GGCGGCAAGCCTGGCTATGCCCTGTTCCAGCTTAGCGATATAGCGGCGCACCTGG<br>GCCGCGCCCCACTCCCGGCGCGTGTAGCGGATGATGCCGCGTAGATCGGCTTCGG<br>CCTCAGCCGTGAGGATGTAGGCCGTCAAGCGCGATCCCCGCTGAGTTCTTCATCA<br>AGAATTTCGCCGACGCTCTTGGTGGACACCTTGCCGGCAAGCCCATCGTTGATGC<br>GGTTCCCCAGCATGGTTTTCAGTTCCTGCCATGCCTGATCGGCATCAGCGTCACCG<br>GGGAACAGACGTTCGAGGGCGTATTGCTTAATGGTCTTGCCCTGCAAGGCGGCCA<br>GGGCTTTCAGGCTCTGGTGCTGCTGGTCCGTCATGTCGATTGTCAGGCGGCTCATT<br>GGATAACCTCCATAAAATACACGTAACCACATTAGCACATATGTGGGCGTGAGGC<br>TACAGCGCGAGGCGCATTAAGGTCGGGAAAATGCGCTAGGCGCATTTAAATTGCG<br>TATTGCTGTAATGCGCCATGCCGGCTAGACTAGGCCCAAATGGGTATACCCAATTT<br>GACCAAGGGGGACGCGATGAGGGCGGCCAAGCACTACCGACAACTTCTATCCAT<br>CGACTTCAACATCGAGGCGCTGGCCTTCGTGCCTGGACCCGACGGCACACGCGGC<br>CGGCGCATCCACGTCCTGGGGCGCGAGGTCCGCGACCGGCCCGGCCTGGTCGAGT<br>ACCTTTCGCCGGCGTTCGGCTCGCGGGTGGCGCTGGACGGCTACTGCAAGGCCAA<br>TTTCGATGCAGTGCTGCACCTGGCGTACCCCGATCATCAGCAATGGGGCCACGCA<br>TGAAGCGCCGAAGCTACGCCATGCTGCGCGCCGCTGCCGCGCTGGCCGTCCTGGT |

CGTTGCCTCGCCGGCATGGGCCGAGCTGCGCGGCGAGGTCGTGCGCATCATCGAC
GGCGACACCATCGACGTGCTGGTAGACAAGCAGCCGGTGCGCGTGCGCCTGGTGG
ACATTGACGCGCCGGAAAAGCGGCAAGCCTTCGGCGAACGTGCGCGCCAGGCGC
TGGCCGGCATGGTGTTCCGCCGGCACGTCCTGGTCGACGAGAAGGACACCGACCG
TTACGGCCGCACGCTGGGCACCGTGTGGGTCAACATGGAGCTGGCCAGCCGGCCG
CCGCAGCCGCGCAACGTCAACGCCGCGATGGTTCACCAGGGCATGGCGTGGGCCT
ATCGCTTCCACGGCCGCGCGGCCGACCCTGAAATGCTGCGGCTCGAACAGGAGGC
GCGAGGCAAGCGCGTCGGCCTCTGGTCCGATCCGCACGCCGTCGAGCCGTGGAAA
TGGCGACGCGAGAGCAACAACCGGAGGGACGAAGGTTGAAGGTCGCCCGCATCT
ACCTGCGCGCCAGTACGGACGAGCAGAATCTTGAACGCCAGGAGAGCCTTGTAGC
GGCCACGCGGGCCGCCGGGTACTACGTCGCCGGCATCTACCGCGAGAAGGCGTCC
GGCGCACGCGCCGACCGGCCCGAGCTGCTGCGCATGATCGCGGACCTGCAACCTG
GTGAAGTCGTCGTTGCGGAGAAGATCGACCGCATCAGCCGCTTGCCGTTGGCCGA
GGCCGAGCGCCTGGTTGCGTCGATCCGGGCCAAAGGGGCCAAGCTGGCCGTGCCT
GGCGTGGTGGACCTGTCGGAGCTGGCCGCCGAGGCGAACGGAGTGGCGAAAATC
GTTCTGGAATCCGTCCAGGACATGCTTTTGAAGCTCGCCTTGCAGATGGCCCGCGA
CGACTACGAGGATCGGCGCGAGCGTCAACGTCAGGGTGTCCAGTTGGCGAAGGC
CGCCGGCCGCTACACCGGCCGCAAACGTGACGCCGGCATGCACGACCGCATCATC
ACGCTTCGCTCCGGCGGATCGAGCATTGCCAAGACGGCCAAGCTGGTCGGATGCA
GCCCGAGCCAGGTCAAACGAGTGTGGGCGGCCTGGAACGCGCAGCAGCAAAAAT
AAAGCCGGGCAGTGCCCGGCTTTTCTCACCTTACCCTAATCTTTCACTTCTATTACT
CAGCTGGCGTTTTTCCATAGGCTCCGCCCCCCTGACGAGCATCACAAAAATCGAC
GCTCAAGTCAGAGGTGGCGAAACCCGACAGGACTATAAAGATACCAGGCGTTTCC
CCCTGGAAGCTCCCTCGTGCGCTCTCCTGTTCCGACCCTGCCGCTTACCGGATACC
TGTCCGCCTTTCTCCCTTCGGGAAGCGTGGCGCTTTCTCATAGCTCACGCTGTAGG
TATCTCAGTTCGGTGTAGGTCGTTCGCTCCAAGCTGGGCTGTGTGCACGAACCCCC
CGTTCAGCCCGACCGCTGCGCCTTATCCGGTAACTATCGTCTTGAGTCCAACCCGG
TAAGACACGACTTATCGCCACTGGCAGCAGCCACTGGTAACAGGATTAGCAGAGC
GAGGTATGTAGGCGGTGCTACAGAGTTCTTGAAGTGGTGGCCTAACTACGGCGAC
ACTAGAAGGACAGTATTTGGTATCTGCGCTCTGCTGAAGCCAGTTACCTTGCGCCA
TTACAACCGGCTATTAGAGTAGCCGGTTTTAGAAAAATTGTCTAAATCGTGATTTT
CCAAATGATTTGAATGATTTGCATGATTGTTTTATACATAAAAAAGAGCGACTCC
TTAATCTCAATTTCGTTTAAGGAATCGCTCACCCAAATATATATCTTTATGTATATT
TAAATATCGTTTAATATCTAAATATACAAGATTATAAAAACAACTCAGTGTTTTTT
TCTTTGAATGATGTCGTTCTGCAACTTTGGCGAGGGACAGAGCGACTCCTTTTTAT
TTTGTTGTCGGCTAGCATTGTACCTAGGACTGAGCTAGCCGTAAAGAAGAGGTTG
GTGATGATAAGATATTTGGCGACCAACGCGGCCTTTTTACGGTTCCTGGCCTTTTG
CTGGCCTTTTGCTCACATGTTCTTTCCTGCGTTATCCCCTGATTCTGTGGATAACCG
TATTACCGCCTTTGAGTGAGCTGATACCGCTCGCCGCAGCCGAACGACCGAGCGC
AGCGAGTCAGTGAGCGAGGAAGCCTGCAGCTCGAGTTAGGATCCTATAAACGCA
GAAAGGCCCACCCGAAGGTGAGCCAGTGTGACTCTAGTAGAGAGCGTTCACCGA
CAAACAACAGATAAAACGAAAGGCCCAGTCTTTCGACTGAGCCTTTCGTTTTATTT
GATGCCTGGAGATCCTTATTAAGCACCGGTGGAGTGACGACCTTCAGCACGTTCG
TACTGTTCAACGATGGTGTAGTCTTCGTTGTGGGAGGTGATGTCCAGTTTGATGTC
GGTTTTGTAAGCACCCGGCAGCTGAACCGGTTTTTTAGCCATGTAGGTGGTTTTAA
CTTCAGCGTCGTAGTGACCACCGTCTTTCAGTTTCAGACGCATTTTGATTTCACCTT
TCAGAGCACCGTCTTCCGGGTACATACGTTCGGTGGAAGCTTCCCAACCCATGGTT
TTTTTCTGCATAACCGGACCGTCGGACGGGAAGTTGGTACCACGCAGTTTAACTTT
GTAGATGAACTCACCGTCTTGCAGGGAGGAGTCCTGGGTAACGGTAACAACACCA
CCGTCTTCGAAGTTCATAACACGTTCCCATTTGAAACCTTCCGGGAAGGACAGTTT
CAGGTAGTCCGGGATGTCAGCCGGGTGTTTAACGTAAGCTTTGGAACCGTACTGG
AACTGCGGGGACAGGATGTCCCAAGCGAACGGCAGCGGACCACCTTTGGTAACTT
TCAGTTTAGCGGTCTGGGTACCTTCGTACGGACGACCTTCACCTTCACCTTCGATT
TCGAACTCGTGACCGTTAACGGAACCTTCCATACGAACTTTGAAACGCATGAACT
CTTTGATAACGTCTTCGCTACTCGCCATAGATCCTTTCTCCTCTTTCAGATCCGTGC

TCAGTATCTCTATCACTGATAGGGATGTCAATCTCTATCACTGATAGGGAAGATCT
CATGAATTCCAGAAATCATCCTTAGCGAAAGCTAAGGATTTTTTTTATCTGAAATT
CTGCCTCGTGATACGCCTATTTTTATAGGTTAATGTCATGATAATAATGGTTTCTTA
GACGTCAGGTGGCACTTTTCGGGGAAATGTG

**Table S3. The sequences of promoters used in this study.**

| Promoters | Sequence (5' to 3') |
|---|---|
| J23119 | TTGACAGCTAGCTCAGTCCTAGGTATAATACTAGT |
| J23101 | TTTACAGCTAGCTCAGTCCTAGGTATTATGCTAGC |
| J23110 | TTTACGGCTAGCTCAGTCCTAGGTACAATGCTAGC |
| J23115 | TTTATAGCTAGCTCAGCCCTTGGTACAATGCTAGC |
| Plac | AAAAAATTTATCAAAAAGAGTGTTGACTTGTGAGCGGATAACAATGATACTTAG ATTCAATTGTGAGCGGATAACAATTTCACACA |
| Pcym | GAAAACAAACAGACAATCTGGTCTGTTTGTATACAGGAAAATTTTTCTGTATAA TAGATTCAACAAACAGACAATCTGGTCTGTTTGTATTAT |

**Table S4. The sequences of riboswitches used in this study.**

| Riboswitches | Sequence (5' to 3') |
|---|---|
| Cbe (Z-inducible) | ATATTAGATATTAGTCATATGACTGACGGAAGTGGAGTTACCACATGAAGTATGACTAGGCATATTATCTTATATGCCACAAAAAGCCGACCGTCTGGGCAAAAAAAGCCTGGATTGCGTCGGCTTTTTTATATGGAAA |
| yxjA (2AP-inducible) | ATCTTAGAAAAAGACATTCTTGTATATGATCAGTAATATGGTCTGATTGTTTCTACCTAGTAACCGTAAAAAACTAGATTACAAGAAAGTTTGAATAAATTTGAACGAGTTGAAAAGGACAAGTTCTTTTCTGTTTGCTCTTATTTTTCACACTTTCTGCACTTCCAGAATTTGTGAAGGATAAGAGCTTTTTTTGTTTCCATAATAACCCTCATAGGAGTTGCTATC |

**Table S5. The sequences of pT181 regulatory elements used in this study.**

| pT181 Regulatory Elements | Sequence (5' to 3') |
|---|---|
| WT-target | AACAAAATAAAAAGGAGTCGCTCACGCCCTGACCAAAGTTTGTGAACGACAT CATTCAAAGAAAAAAACACTGAGTTGTTTTTATAATCTTGTATATTTAGATAT TAAACGATATTTAAATATACATAAAGATATATATTTGGGTGAGCGATTCCTTA AACGAAATTGAGATTAAGGAGTCGCTCTTTTTTATGTATAAAAACAATCATG CAAATCATTCAAATCATTTGGAAAATCACGATTTAGACAATTTTTCTAAAACC GGCTACTCTAATAGCCGGTTGTAA |
| WT-repressor | ATACAAGATTATAAAAACAACTCAGTGTTTTTTTCTTTGAATGATGTCGTTCA CAAACTTTGGTCAGGGCGTGAGCGACTCCTTTTTATTT |
| LS-target | AACAAAATAAAAAGGAGTCGCTCTGTCCCTCGCCAAAGTTGCAGAACGACAT CATTCAAAGAAAAAAACACTGAGTTGTTTTTATAATCTTGTATATTTAGATAT TAAACGATATTTAAATATACATAAAGATATATATTTGGGTGAGCGATTCCTTA AACGAAATTGAGATTAAGGAGTCGCTCTTTTTTATGTATAAAAACAATCATG CAAATCATTCAAATCATTTGGAAAATCACGATTTAGACAATTTTTCTAAAACC GGCTACTCTAATAGCCGGTTGTAA |
| LS-repressor | ATACAAGATTATAAAAACAACTCAGTGTTTTTTTCTTTGAATGATGTCGTTCT GCAACTTTGGCGAGGGACAGAGCGACTCCTTTTTATTT |
| YS-target | AACAAAATAAAAAGGAGTCGCTCGTACCCTCTGCAAAGTTAACGAACGACAT CATTCAAAGAAAAAAACACTGAGTTGTTTTTATAATCTTGTATATTTAGATAT TAAACGATATTTAAATATACATAAAGATATATATTTGGGTGAGCGATTCCTTA AACGAAATTGAGATTAAGGAGTCGCTCTTTTTTATGTATAAAAACAATCATG CAAATCATTCAAATCATTTGGAAAATCACGATTTAGACAATTTTTCTAAAACC GGCTACTCTAATAGCCGGTTGTAA |
| YS-repressor | ATACAAGATTATAAAAACAACTCAGTGTTTTTTTCTTTGAATGATGTCGTTCG TTAACTTTGCAGAGGGTACGAGCGACTCCTTTTTATTT |
| F6-target | AACAAAATAAAAAGGAGTCGCTCACTTACGAACTTGGCGGAACGACGTGAA CGACATCATTCAAAGAAAAAAACACTGAGTTGTTTTTATAATCTTGTATATTT AGATATTAAACGATATTTAAATATACATAAAGATATATATTTGGGTGAGCGA TTCCTTAAACGAAATTGAGATTAAGGAGTCGCTCTTTTTTATGTATAAAAACA ATCATGCAAATCATTCAAATCATTTGGAAAATCACGATTTAGACAATTTTTCT AAAACCGGCTACTCTAATAGCCGGTTGTAA |
| F6-repressor | ATACAAGATTATAAAAACAACTCAGTGTTTTTTTCTTTGAATGATGTCGTTCA CGTCGTTCCGCCAAGTTCGTAAGTGAGCGACTCCTTTTTATTT |
| F4m1-target | AACAAAATAAAAAGGAGTCGCTCACGTTCATGATTGGCGTCAACGATGTGAA CGACATCATTCAAAAGAAAAAAACACTGAGTTGTTTTTATAATCTTGTATATT TAGATATTAAACGATATTTAAATATACATAAAGATATATATTTGGGTGAGCG ATTCCTTAAACGAAATTGAGATTAAGGAGTCGCTCTTTTTTATGTATAAAAAC AATCATGCAAATCATTCAAATCATTTGGAAAATCACGATTTAGACAATTTTTC TAAAACCGGCTACTCTAATAGCCGGTTGTAA |
| F4m1-repressor | ATACAAGATTATAAAAACAACTCAGTGTTTTTTTCTTTGAATGATGTCGTTCA CATCGTTGACGCCAATCATGAACGTGAGCGACTCCTTTTTATTT |

| | |
|---|---|
| F4-target | AACAAAATAAAAAGGAGTCGCTCACGTTCAACTTTGGCGAGTACGATGTGAA<br>CGACATCATTCAAAGAAAAAAACACTGAGTTGTTTTTATAATCTTGTATATTT<br>AGATATTAAACGATATTTAAATATACATAAAGATATATATTTGGGTGAGCGA<br>TTCCTTAAACGAAATTGAGATTAAGGAGTCGCTCTTTTTTATGTATAAAAACA<br>ATCATGCAAATCATTCAAATCATTTGGAAAATCACGATTTAGACAATTTTTCT<br>AAAACCGGCTACTCTAATAGCCGGTTGTAA |
| F4-repressor | ATACAAGATTATAAAAACAACTCAGTGTTTTTTTCTTTGAATGATGTCGTTCA<br>CATCGTACTCGCCAAAGTTGAACGTGAGCGACTCCTTTTTATTT |
| F3m1-target | AACAAAATAAAAAGGAGTCGCTCACGCCTCGAAGTTGGCGCAACGCAGTGT<br>GAACGACATCATTCAAAGAAAAAAACACTGAGTTGTTTTTATAATCTTGTAT<br>ATTTAGATATTAAACGATATTTAAATATACATAAAGATATATATTTGGGTGA<br>GCGATTCCTTAAACGAAATTGAGATTAAGGAGTCGCTCTTTTTTATGTATAAA<br>AACAATCATGCAAATCATTCAAATCATTTGGAAAATCACGATTTAGACAATT<br>TTTCTAAAACCGGCTACTCTAATAGCCGGTTGTAA |
| F3m1-repressor | ATACAAGATTATAAAAACAACTCAGTGTTTTTTTCTTTGAATGATGTCGTTCA<br>CACTGCGTTGCGCCAACTTCGAGGCGTGAGCGACTCCTTTTTATTT |
| F15m5-target | CGACAACAAAATAAAAAGGAGTCGCTCATCTGATTATTGATTTCGGGGGAAA<br>CCATTTAATCATATGAACGACATCATTCAAAGAAAAAAACACTGAGTTGTTT<br>TTATAATCTTGTATATTTAGATATTAAACGATATTTAAATATACATAAAGATA<br>TATATTTGGGTGAGCGATTCCTTAAACGAAATTGAGATTAAGGAGTCGCTCTT<br>TTTTATGTATAAAAACAATCATGCAAATCATTCAAATCATTTGGAAAATCAC<br>GATTTAGACAATTTTTCTAAAACCGGCTACTCTAATAGCCGGTTGTAA |
| F15m5-repressor | CCCCGAAATCAATAATCAGATGAGCGACTCCTTTTTATTT |

**Table S6. Examples of DNA fragments used to create the plasmid library with priming RNA truncations.** Forward amplification primer-Linker-BsmBI site-RNA Primer-BsmBI site-Reverse amplification primer.

| Truncation | Sequence (5' to 3') |
|---|---|
| - 0 nt | CAGATATCGATTCGTGCCAGGTCGTCTCAGCAAACAAAAAAACCACCGCTACCA GCGGTGGTTTGTTTGCCGGATCAAGAGCTACCAACTCTTTTTCCGAAGGTAACTG GCTTCAGCAGAGCGCAGATACCAAATACTGTCCTTCTGCCGTAGTTAGGCCACC AGAACTCTGAGACGTGGAGCTAATTCATCTCGAGGC |
| - 1 nt | CAGATATCGATTCGTGCCAGCGTCGTCTCAGCAACAAAAAAACCACCGCTACCA GCGGTGGTTTGTTTGCCGGATCAAGAGCTACCAACTCTTTTTCCGAAGGTAACTG GCTTCAGCAGAGCGCAGATACCAAATACTGTCCTTCTGCCGTAGTTAGGCCACC AGAACTCTGAGACGTGGAGCTAATTCATCTCGAGGC |
| - 98 nt | CAGATATCGATTCGTGCCAGCATATCGGTCGTTCGTTGCGACTGTCAGCTGGAGT AAGCCGAGTAGTGTTTAATGGCATCATCAATTTGGCACCTGTCAACATATGCCA CGCCATACGGTCGTCTCAGCAAAAATACTGTCCTTCTGCCGTAGTTAGGCCACC AGAACTCTGAGACGTGGAGCTAATTCATCTCGAGGC |
| - 99 nt | CAGATATCGATTCGTGCCAGCATATCGGTCGTTCGTTGCGACTGTCAGCTGGAGT AAGCCGAGTAGTGTTTAATGGCATCATCAATTTGGCACCTGTCAACATATGCCA CGCCATACGAGTCGTCTCAGCAAAATACTGTCCTTCTGCCGTAGTTAGGCCACC AGAACTCTGAGACGTGGAGCTAATTCATCTCGAGGC |

**Table S7. Golden Gate assembly primers used to replace the regulatory elements in pSynORI.** BsmBI site: CGTCTC, Overhang: CGAC, Promoter annealing region: NN…NN.

| Primer's purpose | Sequence (5' to 3') |
|---|---|
| **pT181 target replacement** | |
| Target-Forward | GTCGTCTCACGACAACAAAATAAAAAGGAGTCGCTC |
| Target-Reverse | GTCGTCTCAGCCATTACAACCGGCTATTAGAGTAG |
| Backbone-Forward | CACGTCTCAGTCGGCTAGCATTGTACCTAGG |
| Backbone-Reverse | GTCGTCTCATGGCGCAAACAAAAAAACCAC |
| **pT181 repressor replacement** | |
| Repressor-Forward | GTCGTCTCATGCGAGTTCACCGACAAACAAC |
| Repressor-Reverse | CACGTCTCACGACATACAAGATTATAAAAACAACTCAGTG |
| Backbone-Forward | CACGTCTCACGCACTTAACATCAATCTAATTATATATCATTATTAC |
| Backbone-Reverse | GTCGTCTCAGTCGACTAGTATTATACCTAGGACTGAG |
| **pT181 target promoter replacement** | |
| Target-Forward | GTCGTCTCACGACAACAAAATAAAAAGGAGTCGCTC |
| Target-Reverse | GTCGTCTCAGCCATTACAACCGGCTATTAGAGTAG |
| Backbone-Forward | CACGTCTCAGAAGAGGTTGGTGATGATAAGATATTTG |
| Backbone-Reverse | CACGTCTCATGGCGCAAAGGTAACTGG |
| Promoter-Forward | CTTCNN…NN |
| Promoter-Reverse | GTCGNN…NN (complementary sequence) |
| **pT181 repressor promoter replacement** | |
| Repressor-Forward | GTCGTCTCATGCGAGTTCACCGACAAACAAC |
| Repressor-Reverse | CACGTCTCACGACATACAAGATTATAAAAACAACTCAGTG |
| Backbone-Forward | GTCGTCTCAGAAGACTCAATACCTACCCATTCC |
| Backbone-Reverse | CACGTCTCACGCACTTAACATCAATCTAATTATATATCATTATTAC |
| Promoter-Forward | CTTCNN…NN |
| Promoter-Reverse | GTCGNN…NN (complementary sequence) |

**Movie S1.**

Movie of cells with IPTG-inducible pSynORI, after cells were loaded, fresh LB media (20 mL in 60 mL syringe) with antibiotic and 0.5% Tween 20 were added to the channel with a flow velocity of 100 μm/s through the channel. After cells fulfilled the trap and stabilized, the channel's input media was switched to fresh LB media (20 mL in 60 mL syringe) with 100 μM IPTG, 10 μL Oregon Green 488 dye (Fisher), corresponding antibiotic, and 0.5% Tween 20 with the same flow velocity. After 17 hours, the input media was switched back to LB media without IPTG till the end of the experiment.

**Movie S2.**

Movie of cells with both IPTG-inducible and cumate-inducible pSynORIs, after cells were loaded, fresh LB media (20 mL in 60 mL syringe) with 100 μM IPTG, antibiotics and 0.5% Tween 20 were added to the channel with a flow velocity of 50 μm/s through the channel. After cells fulfilled the trap and stabilized, the channel's input media was switched to fresh LB media (20 mL in 60 mL syringe) with 200 μM cumate, corresponding antibiotics, and 0.5% Tween 20 with the same flow velocity. After 17.8 hours, the input media was switched back to LB media with 100 μM IPTG till the end of the experiment. Both types of media contained no dye, and the media switch time was recorded manually during the switch.
